# Simultaneous EEG-PET-MRI identifies temporally coupled, spatially structured hemodynamic and metabolic dynamics across wakefulness and NREM sleep

**DOI:** 10.1101/2025.01.17.633689

**Authors:** Jingyuan E. Chen, Laura D. Lewis, Sean E. Coursey, Ciprian Catana, Jonathan R. Polimeni, Jiawen Fan, Kyle S Droppa, Rudra Patel, Hsiao-Ying Wey, Catie Chang, Dara S. Manoach, Julie C. Price, Christin Y. Sander, Bruce R. Rosen

## Abstract

Sleep entails significant changes in cerebral hemodynamics and metabolism. Yet, the way these processes evolve throughout wakefulness and sleep and their spatiotemporal dependence remain largely unknown. Here, by integrating a novel functional PET technique with simultaneous EEG-fMRI, we reveal a tightly coupled temporal progression of global hemodynamics and metabolism during the descent into NREM sleep, with large hemodynamic fluctuations emerging as global glucose metabolism declines, both of which track EEG arousal dynamics. Furthermore, we identify two distinct network patterns that emerge during NREM sleep: an oscillating, high-metabolism sensorimotor network remains active and dynamic, whereas hemodynamic and metabolic activity in the default-mode network is suppressed. These results elucidate how sleep diminishes awareness while preserving sensory responses, and uncover a complex, alternating balance of neuronal, hemodynamic, and metabolic dynamics in the sleeping brain. This work also demonstrates the potential of EEG-PET-MRI to explore neuro-hemo-metabolic dynamics underlying cognition and arousal in humans.

## Introduction

During sleep, neuronal, hemodynamic, and metabolic dynamics undergo profound changes to accommodate diverse functions of this complex, active, and restorative state. For instance, mounting evidence has demonstrated a substantial change in cerebral hemodynamics during sleep^1–4^, which not only reflects marked changes in neuronal oscillations and autonomic processes^5–8^, but also drives cerebrospinal fluid (CSF) dynamics linked to metabolic waste clearance^9,10^. Additionally, the brain must maintain a delicate balance in which energy consumption is reduced to enter the sleeping state, yet with sufficient remaining metabolic activity to enable sensory-alerting or awakening when needed^11^. To date, how these metabolic dynamics evolve throughout the brain across arousal states, and their spatiotemporal coordination with brainwide hemodynamic waves remain unclear. Addressing these questions will not only deepen our understanding of sleep physiology, elucidating the complex neuro-hemo-metabolic balance underlying the gradual descent into sleep while facilitating sensory-driven awakening, but can also shed new light on disease mechanisms. For instance, how the balance of neuronal waste production (metabolism) and clearance (CSF changes driven by hemodynamics) is disturbed in various sleep disorders, contributing to neurodegeneration and neuroinflammation^12,13^.

A major hurdle in investigating the time-varying evolution of hemodynamic and metabolic processes in humans has been a lack of multimodal imaging tools to directly map and link their temporal dynamics during sleep. While functional magnetic resonance imaging (fMRI) enables high-resolution, whole-brain measurements of functional and physiological dynamics in sleep, it cannot provide direct energetic information: blood-oxygenation-level-dependent (BOLD) fMRI signals and metabolism can dissociate in many settings^14–16^. MR spectroscopy-based metabolic imaging is constrained by limited spatial resolution, sensitivity, and brain coverage^17^. Additionally, while traditional [¹⁸F]Fluorodeoxyglucose (FDG)-based positron emission tomography (PET) can quantify glucose metabolism and has been previously employed to characterize the differential metabolic patterns between wake and different phases of sleep^18–21^, it has classically been limited to slow timescales that could not access dynamic metabolic information over the course of a scan.

Recent advances in functional PET (fPET)-FDG imaging have provided an opportunity for tracking rapid dynamics of glucose uptake non-invasively in humans, through the introduction of a constant tracer infusion paradigm^22^; and emerging evidence has demonstrated its potential for imaging metabolic dynamics at timescales approaching one minute or below^23,24^. Additionally, since fPET-FDG is carried out in one imaging session, circumventing the inter-scan variability faced by conventional PET paradigms, it is expected to have superior imaging sensitivity. Thus, fPET-FDG holds the promise of revealing more rapid and more subtle variations of glucose uptake in sleep that approach the spatiotemporal scales of fMRI-based hemodynamic measures. With the integrated PET-MR scanners, both measures can now be made simultaneously.

Here, we integrate fPET-FDG with simultaneous EEG-fMRI, aiming to elucidate the spatiotemporal dependence of the brain’s rich hemodynamics and metabolic dynamics across wakefulness and sleep. By introducing a time-resolved framework that unifies dynamic information across timescales, we show that global hemodynamic and metabolic changes are temporally coupled across wakefulness and sleep. Specifically, increased global fMRI fluctuations (on timescales of seconds, locked to low-frequency neuronal, autonomic, and CSF fluctuations^8–10^) coincide with reduced glucose uptake (on timescales of minutes), implying temporally coordinated neuronal, vascular, and metabolic dynamics accompanying arousal state fluctuations. Furthermore, we identify two distinct hemo-metabolic patterns that emerge during NREM sleep: sensory networks exhibit high-amplitude, low-frequency (peaking at ∼0.02 Hz) hemodynamic waves and relatively preserved metabolic activity, while the default-mode network^25^ exhibits smaller hemodynamic fluctuations and the most pronounced metabolic decline. These observations reveal spatially distinct effects of sleep, supporting a pathway that may potentially facilitate sensory-driven alerting and awakening while suppressing higher-order cognition during sleep. From a methodological perspective, the multi-modal imaging and analytical framework established in this study also enables us to illuminate how neuronal, hemodynamic, and metabolic dynamics interact to mediate cognition and consciousness, at a temporal resolution not feasible previously in humans.

## Results

### Tracking electrophysiological, hemodynamic and metabolic dynamics across sleep-wake cycles using tri-modal imaging

We integrated functional PET with simultaneous EEG-fMRI to achieve joint measures of time-varying changes in electrophysiology, hemodynamic signals, and glucose uptake accompanying arousal state transitions. Concurrent EEG recordings (*N*=21 subjects) or behavioral data (*N*=5 subjects) were collected to infer the dynamic sleep or wakefulness state of the subjects in the scanner. See ***Methods*** for the overall design of the experiment and data acquisition. The statistics of sleep-wake durations and the histogram of arousal state variations over time are summarized in the supplementary Fig. S1.

Our tri-modal imaging framework captured arousal-induced dynamic variations of electrophysiological, hemodynamic, and metabolic signals (Fig. 1). Consistent with previous studies^26,27^, we observed a prominent increase in the amplitudes of fMRI signal fluctuations in NREM sleep; and a marked reduction of glucose uptake was identified, evidenced by decreased slopes in fPET time-activity curves (TACs). Of note, in addition to the significant metabolic differences between wake and NREM sleep, fPET-FDG demonstrated sufficient sensitivity to capture more subtle variations of metabolic patterns within NREM sleep (Fig. 1B).

**Figure 1:**
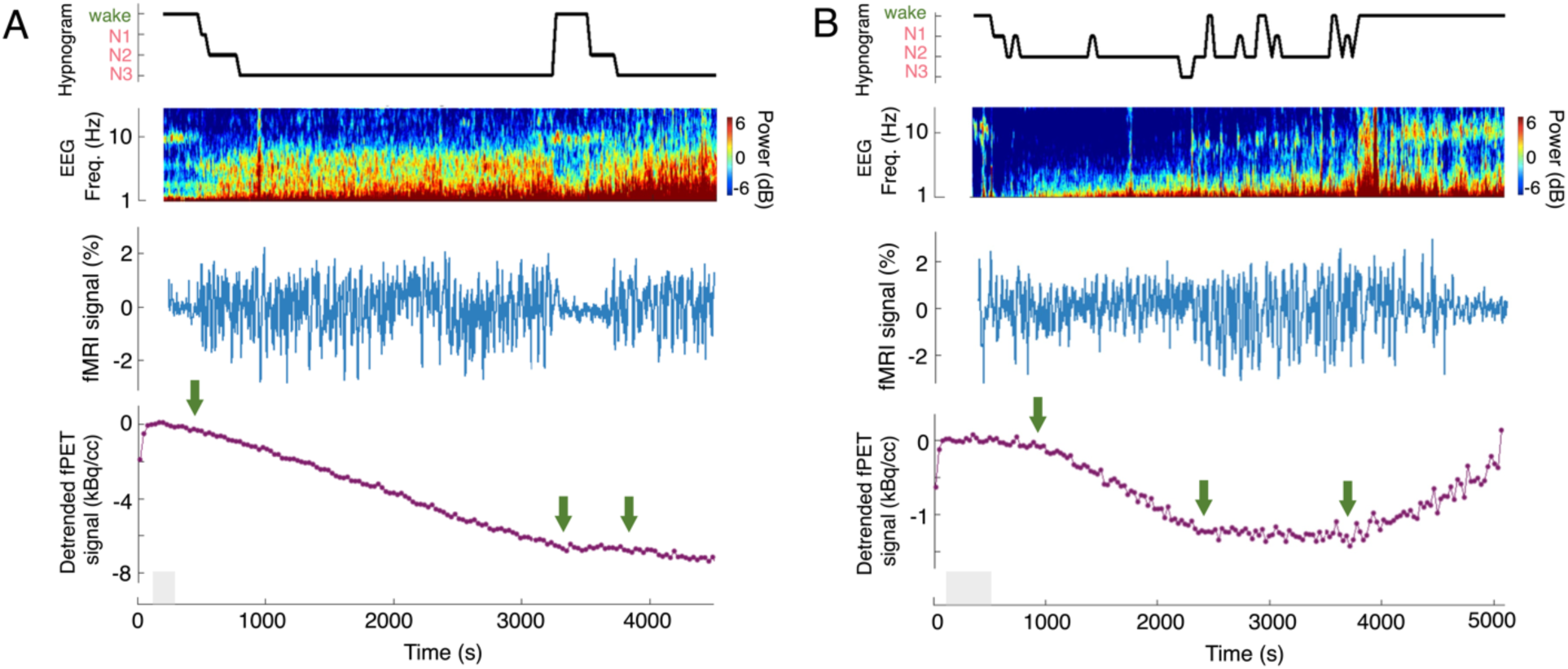
Tri-modal imaging of the electrophysiological, BOLD-fMRI, and fPET-FDG metabolic dynamics accompanying arousal-state transitions. **(A, B)** Top: Hypnogram of scored sleep staging and the spectrogram of an occipital EEG electrode; middle: fMRI-based hemodynamic oscillations of the visual network; bottom: fPET-based metabolic dynamics of the default-mode network. Networks were extracted using a public functional atlas^52^. Functional PET signals were temporally detrended according to the arbitrarily chosen initial wakeful period (i.e., removal of the linear trend fitted to the data points within the shaded gray area) only in this plot to help visualize altered slopes at state transitions, with an increase/decrease of TAC slope indicating increased/decreased metabolism. Changes in electrophysiological recordings, fMRI intensities, and glucose metabolism (highlighted with green arrows) were identifiable across sleep-wake cycles (**A**, subject 1) and within the NREM sleep (**B**, subject 2), mirroring arousal-state transitions (top, inferred from simultaneous EEG recordings). Note that our goal here is to highlight arousal-induced changes in the imaging signals, so we show fMRI and fPET signals from exemplar networks that exhibit strong sleep-wake differences for each modality independently (see group-level results in Fig. 3 below; to avoid a circular analysis, we re-ran the group-level analysis without including the two illustrative subjects shown here, the findings remained unaltered).

### Temporally coupled hemodynamic and metabolic changes across wakefulness and NREM sleep

Having shown the sensitivity of our multi-modal method, we first examined the temporal association between time-varying metabolic and hemodynamic patterns over the sleep-wake cycle. During NREM sleep, the amplitude of fMRI fluctuations in the 0.01-0.1 Hz range strongly increases^27^, and these fluctuations correspond to low-frequency oscillations in neuronal and autonomic dynamics^4,8^, which further drives CSF dynamics^9,10^. However, the metabolic state corresponding to these low-frequency oscillations is not known. We quantified the amplitude of fMRI fluctuations and linked it to the simultaneously measured slow-scale changes in glucose uptake over time. To integrate the distinct timescales of fPET and fMRI signals, we calculated the integral of time-windowed measures of the amplitude variation of BOLD fluctuations (BOLD-AV, see ***Methods***), and tested its correlation with the temporally detrended fPET-FDG TACs to assess the temporal dependence of observed hemodynamic and metabolic changes. We refer to these “quasi-metabolic” TACs, modeled by BOLD-AV and sharing similar temporal properties with fPET-FDG signals, as BOLD-AV-TACs. This time-resolved scheme of fPET-fMRI integration (illustrated in Fig. 2A, and see Fig. S2 for individual fitting results) was motivated by the accumulative nature of PET-FDG tracer kinetics and built upon the construction of stimulus-driven metabolic regressor in existing fPET studies, i.e., treating BOLD-AV as an indicator of the power of instantaneous functional activity of interest here. Cross-correlation analyses revealed a tight temporal coupling between the global BOLD-AV-TACs and fPET-FDG TACs (Fig. 2B), suggesting coordinated hemodynamic changes and glucose metabolism across wake and NREM sleep. The temporal dependence of hemodynamic and metabolic signals was further corroborated by a linear regression analysis: using the global BOLD-AV-TAC as the covariate to regress against brain-wide fPET-FDG data revealed several cortical regions that demonstrated strong metabolic covariations with BOLD-AV-TAC (Fig. 2C). Such fPET-fMRI temporal association was also identifiable at region-specific scales, see supplementary Fig. S3 for the regression results using region-specific BOLD-AV-TAC. These results demonstrated a coupled evolution of cerebral dynamics in NREM sleep, with large, seconds-scale hemodynamic fluctuations co-occurring with a drop in metabolic rate over minutes.

**Figure 2:**
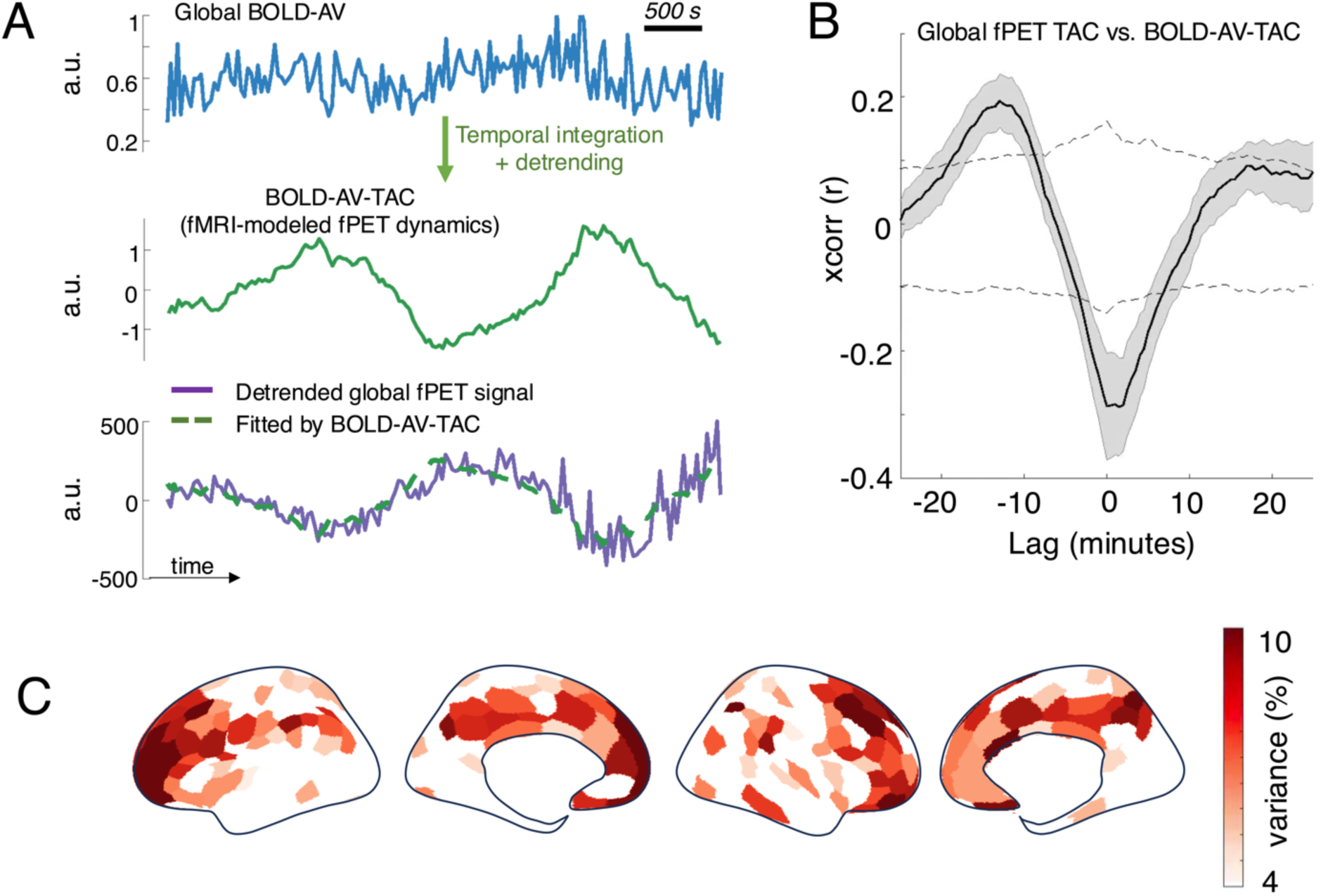
Temporal coupling between the global hemodynamic and metabolic changes across wakefulness and sleep. **(A)** Summary of the scheme of linking instantaneous BOLD-AV to fPET-FDG TACs, illustrated using a single subject’s data. To mimic FDG accumulation over time, the amplitude of fMRI fluctuations, BOLD-AV, was temporally integrated for comparison with fPET-FDG TACs. Ultra-slow changes relating to baseline FDG accumulation in BOLD-AV-TAC and fPET-FDG TAC were removed using a third-order polynomial. **(B)** Cross-correlations between the global fPET-FDG TAC and the TAC modeled by the global BOLD-AV (BOLD-AV-TAC), after the removal of baseline trends. Positive lags indicate delayed fPET-FDG TAC relative to BOLD-AV-TAC. Mean and standard errors across subjects are shown. Gray dashed lines indicate 95% confidence intervals of the null cross-correlations, derived nonparametrically from phase-reshuffled data. **(C)** Spatial distributions of the variance of temporally detrended fPET-FDG TACs explained by the global BOLD-AV-TAC (300 ROIs, *N* = 23, FDR, *p* < 0.05). ROI-wise fPET-FDG TACs were temporally smoothed with a median filter using a 2.5-minute (five temporal frames) kernel before variance fitting; identical temporal smoothing was applied to sham data to construct null distributions.

### Spatially structured cortical gray-matter distributions of hemodynamic and metabolic patterns: wake vs. NREM sleep

Having found that hemodynamic oscillations and metabolic declines were tightly coupled in time, we next investigated whether these effects were spatially consistent across the brain. We aimed to understand if these hemodynamic oscillations were a global effect, or signaled network-specific changes in metabolic function. Our simultaneous measures provided an excellent opportunity to investigate the fine-grained spatial relationship between BOLD and metabolic patterns in sleep versus wakefulness, owing to the high-sensitivity BrainPET scanner and fPET protocol. Our simultaneous measures also enabled us to address the substantial across-session inconsistencies of sleep state distributions in independent, sequential fPET and fMRI experiments.

To quantify sleep-induced changes in the amplitudes of fMRI fluctuations, we computed the fractional changes in BOLD-AV during stable NREM sleep compared to the stable wakeful period (see ***Methods***). We found that prominent increases in BOLD-AVs could be identified in almost all brain regions with the most substantial changes in the visual, auditory, and motor cortices (Fig. 3A), at spatial scales consistent with previous sleep fMRI reports^26,27^.

**Figure 3:**
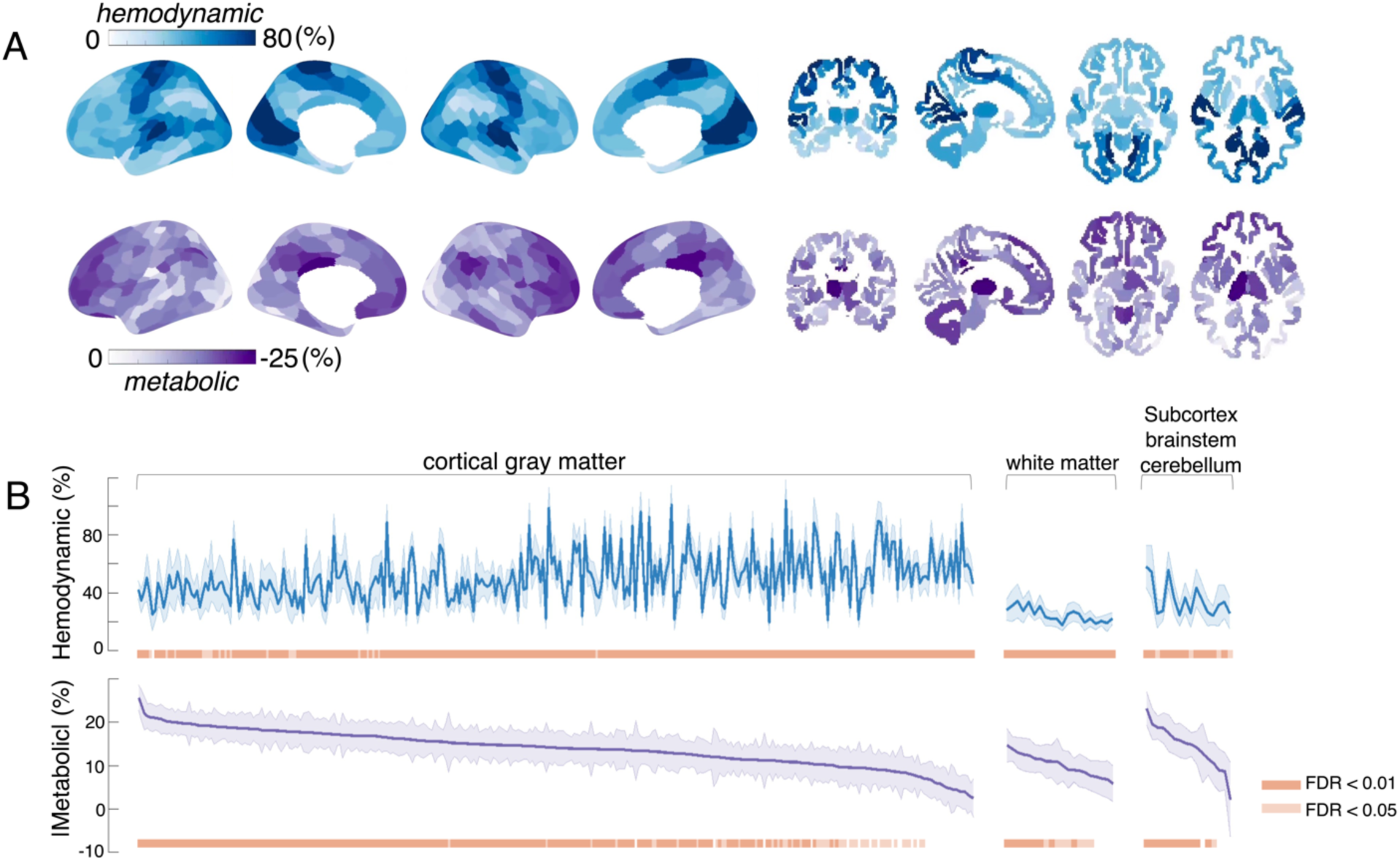
Spatially structured distributions of NREM sleep-induced changes in cortical hemodynamics and glucose metabolism. **(A)** Spatial patterns of fractional increases in BOLD-AV (blue) and decreases in FDG uptake (purple) during NREM sleep compared to wakeful resting, averaged across subjects (300 ROIs, *N*=23). **(B)** ROI-specific quantitative changes in BOLD-AV and FDG uptake (mean and standard errors across subjects). See ***Methods*** for the extraction of cortical, subcortical, and white-matter ROIs. The statistical significance of hemodynamic and metabolic changes of each ROI is shown at the bottom (with two FDR levels). Statistically significant hemodynamic and metabolic changes were identified in many brain regions. ROIs within each group (“cortical gray matter”, “cortical white matter”, and “subcortex/brainstem/cerebellum”) were sorted according to the fractional changes in FDG uptake to help visualize the alignment of sleep effects in BOLD-AV and glucose metabolism.

To measure sleep-induced changes in glucose metabolism, we computed the fractional changes in the slopes of fPET TACs during stable NREM sleep compared to the stable awake period, since in steady state (i.e., approximately constant plasma FDG concentration), the slopes of fPET TACs are proportional to the absolute FDG uptake^22^ (see ***Methods***). We found a widespread drop in cerebral metabolism during NREM sleep (Fig. 3A), which were consistent with early FDG PET studies that quantified sleep vs. wake differences using two separate scans^20,21^, with the frontal cortex, posterior cingulate cortex, and thalamus exhibiting the most salient reductions in glucose uptake during NREM sleep. Notably, owing to the enhanced sensitivity of the BrainPET scanner, we also observed statistically significant metabolic reductions in the brainstem, cerebellum, and white matter.

To test whether the characterized sleep metabolic patterns resulted partially from systematic misspecification of fPET TACs^28^—that the sleep and wakefulness timing distributions interact with the nonlinear baseline of fPET TACs—we performed a control analysis that quantified the group-level fPET-FDG signal slope alterations by applying each subject’s sleep-wake state delineations (see ***Methods***) to the fPET-FDG TACs of two subjects that either stayed awake or asleep for >95% of the time during the experiments (control cases where no prominent arousal-state transition occurred). This control analysis resulted in insignificant sleep-wake differences in glucose uptake for both subjects (Fig. S4), supporting the reliability of characterized sleep metabolic patterns during NREM sleep. Potential bias from fPET baseline misspecification may cancel out at the group level due to varying sleep-wake timings across subjects.

Additionally, to validate the spatial patterns of metabolic changes estimated using direct state-dependent slopes of fPET TACs, we further quantified the FDG uptake using image-derived arterial input functions^29^ and a Patlak graphical analysis^30^, which yielded consistent spatial distributions of metabolic changes as shown in Fig. 3A (Spearman’s spatial correlation = 0.73, *p* < 1e-15, 300 ROIs, see Supplementary Methods, Fig. S6).

Having shown the validity of the measured metabolic changes, we contrasted them to sleep-induced BOLD-AV results. The overall spatial distributions of NREM sleep-induced increases in BOLD fluctuations and metabolic reductions exhibited a significant anti-correlation (Spearman’s spatial correlation = −0.41, *p* = 5.7×1e-14, 300 ROIs). The cortical gray-matter regions demonstrating the strongest increases in BOLD-AV, primary sensory networks, showed the smallest metabolic declines in sleep (see Fig. S7 for sleep-induced BOLD and metabolic changes overlaid on the same cortical surface). These results demonstrated that across the cortex, the regions with large hemodynamic fluctuations also preserved the highest level of metabolic activity during sleep, whereas a suppression of hemodynamic fluctuations indicated a suppression of metabolism. Of note, the anti-correlated relationship between sleep-induced hemodynamic amplitude increases and metabolic reductions were not observed in subcortical regions (Spearman’s spatial correlation = 0.42, *p* = 0.11, 16 ROIs) or across white-matter tracts (Spearman’s spatial correlation = 0.72, *p* = 4.6×1e-4, 22 ROIs) during NREM sleep, which may imply distinct metabolic mechanisms underlying hemodynamic changes beyond cortical gray matter.

### Spatially structured cortical gray-matter distributions of hemodynamic and metabolic patterns: correlation with continuous, gradual changes in EEG arousal

Given that fPET-FDG provides time-resolved measures of glucose uptake, we further compared the spatial signatures of hemodynamic and metabolic changes coupled to continuous arousal fluctuations over time. Having identified differences between discrete wakeful vs. sleep states (Fig. 3), we next assessed how hemodynamic and metabolic levels of different brain regions tracked gradual changes in arousal, including sleep depth variations within the NREM phase.

The instantaneous arousal levels for each subject were inferred from simultaneous EEG measurements, by taking the ratio of signal amplitude in the 7−13 Hz range (alpha wave activity) over the amplitude in the 1−7 Hz range (delta and theta activity)^31^ (see ***Methods*** for the calculation of EEG arousal index, EEG_arousal_, and Fig. 4A for an illustration). As expected, both global hemodynamic and metabolic dynamics were temporally coupled to the EEG arousal index over time (Fig. 4B).

**Figure 4:**
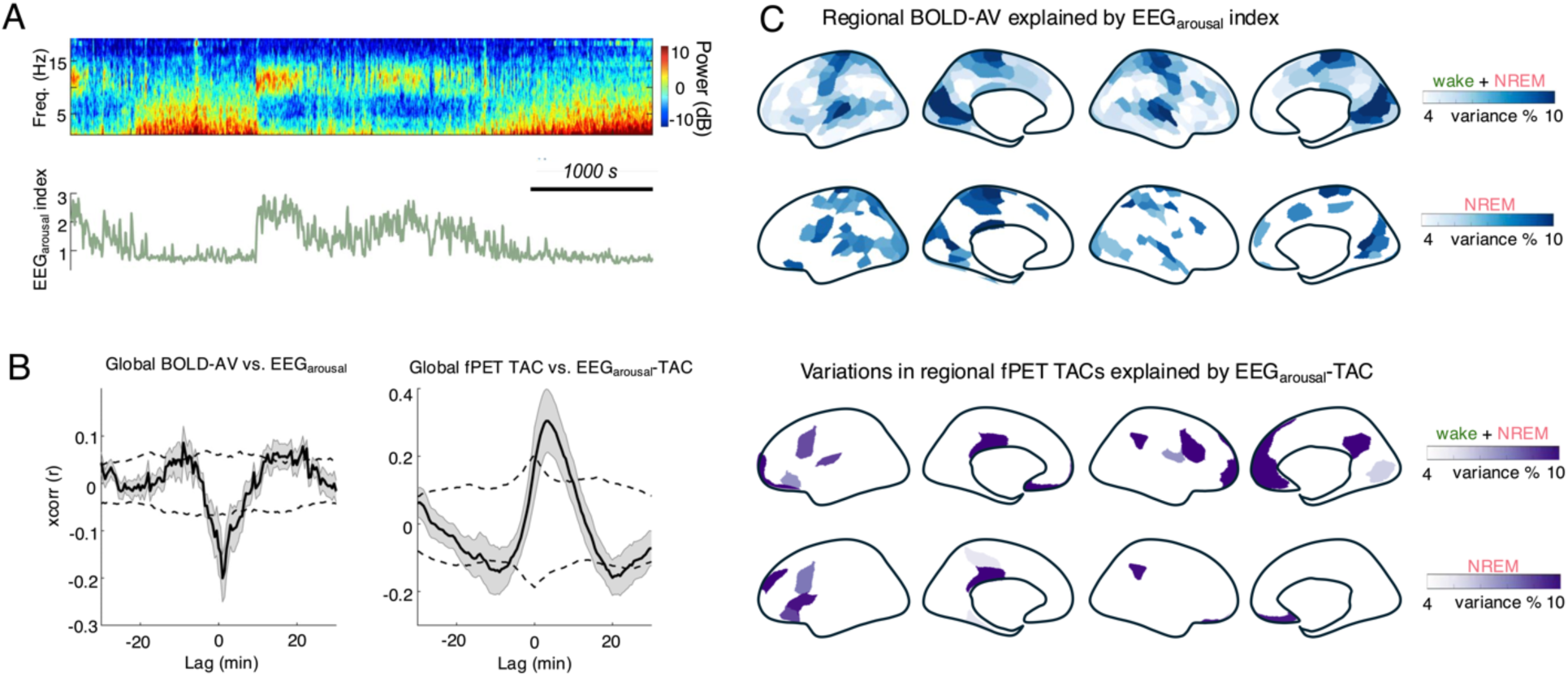
Spatial distributions of cortical hemodynamic and metabolic patterns coupled to gradual changes in EEG arousal. **(A)** Illustration of continuous arousal fluctuations throughout the experiment (“EEG_arousal_ index”) for a single subject, with the spectrogram of an occipital EEG electrode shown at the top for reference. **(B)** Cross-correlation analyses suggested that both the global BOLD-AV and fPET TAC tracked gradual changes in arousal. EEG_arousal_-TAC: TACs modeled by EEG_arousal_ indexes. Positive lags indicate delayed global BOLD-AV/fPET TAC relative to EEG_arousal_/EEG_arousal_-TAC. Mean and standard errors across subjects are shown. Gray dashed lines indicate 95% intervals of the null group-level cross-correlation values, derived nonparametrically from phase-reshuffled data. **(C)** Spatial patterns of hemodynamic and metabolic changes coupled to gradual changes in arousal, across the entire sleep-wake cycle (“wake + NREM”) or during the NREM sleep phase alone (“NREM”). Top: BOLD-AVs explained by EEG_arousal_ under a linear regression model (300 ROIs, FDR, *p*<0.05). Bottom: fPET-FDG TACs explained by EEG_arousal_-TAC (100 ROIs, FDR, *p*<0.05), after temporal detrending. Note that the ROI size was increased to enhance the power for testing fPET-EEG coupling. ROI-wise fPET-FDG TACs were temporally smoothed with a median filter using a 2.5-minute (five temporal frames) kernel before variance fitting; identical temporal smoothing was applied to sham data to construct null distributions. Results were estimated across subjects with clean EEG recordings (*N*=18).

Hemodynamic signatures of gradual arousal modulation were identified by regressing EEG_arousal_ against region-specific BOLD-AV time series. Identical sets of regions, comprising the visual, auditory, and motor cortices, exhibited the strongest associations with arousal fluctuations as those showing the most prominent sleep vs. wake differences (Fig. 4C, top; the Spearman’s spatial correlation between cortical BOLD-AV patterns in Figs. 4C and 3A was 0.88 (*p* < 1e-15, 300 ROIs) for the entire sleep-wake cycle and 0.58 (*p* < 1e-15, 300 ROIs) for the specific NREM sleep phase).

Metabolic signatures of arousal were characterized by regressing the quasi-metabolic TAC modeled by EEG_arousal_ (EEG_arousal_-TAC) against region-specific fPET TACs over time. Amongst the extensive cortical regions that exhibited salient wake vs. sleep FDG metabolic differences, varying effects of arousal state on metabolic dynamics were observed, with the metabolism of the prefrontal and posterior cingulate cortices most deeply suppressed as arousal declined (Fig. 4C, bottom; the Spearman’s spatial correlation between cortical fPET metabolic patterns in Figs. 4C and 3A was 0.72 (*p* < 1e-15, 100 ROIs) for the entire sleep-wake cycle and 0.61 (*p* < 1e-15, 100 ROIs) for the specific NREM sleep phase).

When contrasting fMRI and fPET-derived signatures of EEG-indexed arousal, regions demonstrating the strongest hemodynamic fluctuations vs. the strongest metabolic suppression during gradual variations in arousal and sleep depth were spatially distinct, in line with the wake vs. NREM sleep findings. These results confirmed that a spatially distinct sensory network shows high-amplitude hemodynamic fluctuations and higher metabolic activity, with tight coupling to continuous, dynamic changes in arousal state. By contrast, the default mode network showed smaller hemodynamic oscillations and an overall suppression of metabolic activity.

## Discussion

Arousal state shifts dynamically over time, inducing continuous modulation of the brain’s neurovascular and metabolic function. Here, by integrating simultaneous EEG-fMRI with functional PET-FDG and introducing a time-resolved multi-modal fusion, we uncovered spatiotemporally coordinated cerebral hemodynamic and metabolic processes across different arousal states. Temporally, large fluctuations in hemodynamic signals emerged when the global glucose uptake went down, both tracking EEG arousal fluctuations. Spatially, hemodynamic and metabolic effects were structured in two distinct network patterns locked to arousal, with sensory networks showing greater changes in hemodynamic fluctuations and smaller metabolic alterations accompanying sleep-wake transitions and sleep-depth variations.

The spatial patterns of metabolic changes between wakefulness and NREM sleep are consistent with reductions in energetic costs associated with reduced local neuronal signaling during NREM sleep^11^. Cortices demonstrating the strongest reductions in glucose uptake encompass the set of brain regions commonly referred to as the default-mode network^25^, which are marked by active spontaneous neuronal activity in the wakeful resting state and high metabolic demands in terms of both glucose consumption and anaerobic glycolysis^32^. Within this network, the fPET dynamics of the prefrontal cortex, and the posterior cingulate cortex exhibited the highest sensitivity to graded changes in arousal, consistent with their prominent roles in consciousness and arousal circuitry^33,34^. In contrast to these suppressed default-mode network dynamics, a network comprising sensory cortices exhibited preserved higher metabolic activity, and larger hemodynamic amplitude oscillations in NREM sleep. One possible interpretation is that sensory regions undergo periodic alternations between low and high activity linked to arousability during sleep (as reflected by larger hemodynamic oscillations), the maintenance of which requires higher baseline metabolic rates (as reflected by smaller metabolic reductions than the awake state). We observed that oscillatory hemodynamic waves in NREM sleep peaked at ∼0.02 Hz (Fig. S8A)—consistent with earlier studies, these oscillatory hemodynamic waves coincided with infraslow EEG sigma (spindle) band activity^4^ and cardiac oscillations^8^ in NREM sleep (Figs. S8 and S9, with the strongest correlation between fMRI signals and EEG spindle activity occurring in the sensory network). Coordinated 0.02 Hz oscillations in sigma activity and autonomic processes during NREM sleep have been linked to alternating phases of offline sleep processing and heightened susceptibility to external stimulation^35,36^, with periodic sigma activity coinciding with different levels of sensory arousability^36–38^. Furthermore, reduced metabolism accompanying the emergence of large 0.02 Hz oscillations may be attributable to decreased minute-scale baseline activity during the descent into NREM sleep—similar oscillatory and baseline activity patterns were recently identified in the locus coeruleus^39^, a key regular of arousal that mediates the likelihood of sensory awakening^40,41^. Thus our observations reflect that, during the progression into deep sleep, sensory networks intermittently reach states of heightened arousal, allowing increased pass-through of incoming sensory input, whereas higher-order regions show a more total suppression of neuronal activity. These observations identify a pathway that could facilitate sensory-driven alerting and awakening in NREM sleep.

The observed hemo-metabolic patterns should be considered in the context of the specific sleep phase distributions in our study—while several subjects descended into Stage N3 sleep during the imaging session, our results, especially those encompassing measures across the entire scan, were dominated by the contrast of wakefulness and light and intermediate Stage N1 and N2 sleep (see Fig. S1). We noticed that, in occasional cases where the subject gradually transitioned from N2 to deep N3 sleep, both global and sensory fMRI oscillation amplitudes appeared to decrease when metabolism continued to decline (see supplementary Fig. S10 for an example). Reduced global hemodynamic oscillations in N3 sleep may result from reduced efficiency of the dominating neural rhythms in entraining regional vascular changes despite the increase in global slow wave activity^4,42^, as hemodynamic responses are attenuated when neural events are spaced closely in time. Moreover, sympathetic activity that elicits widespread fMRI responses also decreases in stage N3 than N2 sleep^43^. This pattern likely reflects a nonlinear, inverted-U relationship between vascular oscillations and metabolism, with hemodynamic oscillations arising as slow neural and sympathetic events appear in light/intermediate sleep, and then subsiding as slow waves become denser in deeper sleep, leading to suppressed hemodynamic entrainment and metabolism. Thus degraded temporal correlation between global fMRI and fPET dynamics may be expected in acquisitions that include more deeper stages of sleep. Spatially, reduced sensory hemodynamic oscillations also align with suppressed sensory activity in deep NREM sleep, shedding light on the neuronal mechanisms by which the brain becomes less responsive to sensory stimulation in Stage N3 sleep. Together, our results reveal a complex landscape of network-specific hemodynamic oscillations and baseline metabolic activity across wakefulness and different stages of NREM sleep, as outlined in Fig. 5. Future research encompassing a broader range of the NREM sleep cycle is needed to fully validate the hemo-metabolic patterns unique to N3 sleep.

**Figure 5:**
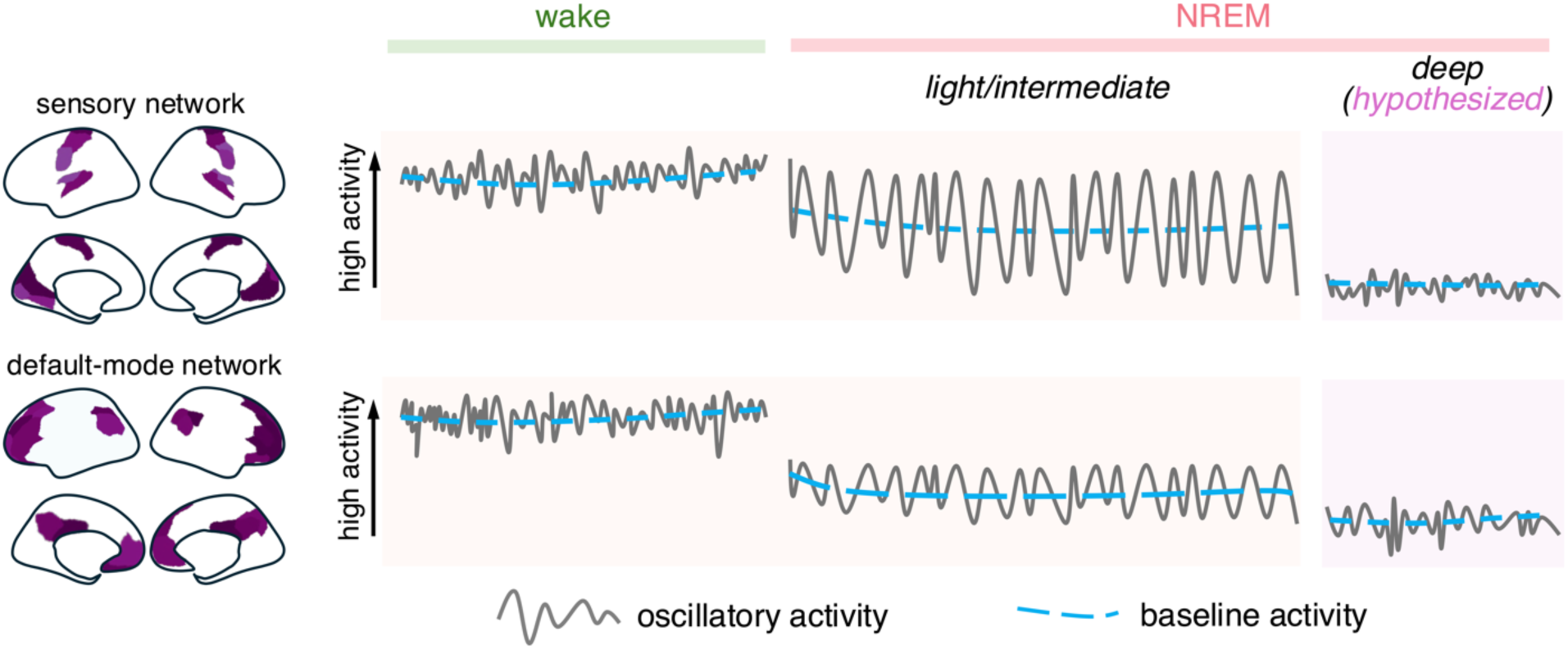
Cartoon illustration of distinct sensory vs. default-mode network activity patterns across wakefulness and different stages of NREM sleep. As the subject transitions from wakefulness to light/intermediate NREM sleep, the sensory network displays prominent ultra-slow oscillatory activity (reflected by large hemodynamic waves) and a modest decline in baseline activity (reflected by reduced metabolism; note that tracking minute-scale baseline activity with fMRI is challenging due to contamination from low-frequency scanner drifts). By contrast, the default-mode network exhibits smaller oscillatory activity and a stronger reduction in baseline activity. Based on observations from a small number of participants undergoing gradual transitions into stable, deeper NREM sleep, we hypothesize that both oscillatory and baseline activity decline across networks as sleep deepens, which requires validation in future studies.

It is of note that while we found that larger hemodynamic fluctuations co-localized with higher metabolic activity in cortical gray matter, the opposite pattern was observed across white-matter tracts (Fig. 3B), where higher hemodynamic fluctuations accompanied stronger decrease in metabolism. This distinction may be attributable to disparate energetic budgets between gray and white matter—instead of signaling, the vast majority of energetic costs in white matter are devoted to non-signaling housekeeping^44^. It is therefore possible that the metabolic and hemodynamic distributions correlated indirectly due to their association with the anatomy of white-matter vasculature: higher vessel densities mirror higher resting metabolic levels, and also result in larger fMRI responses subsequent to a systemic change in blood flow (responding to either global neuronal dynamics or autonomic regulation) during sleep. A recent study provided support for this hypothesis by showing that resting-state white-matter fMRI signal magnitudes scale positively with white-matter microvascular densities^45^.

Regarding the quantitative results of absolute metabolic changes in NREM sleep, it is of note that, in order to increase the sample size of wakeful and sleep periods, we applied a modest threshold of five minutes to identify stable wakeful and sleep segments in each subject’s data (see ***Methods***). Consequently, FDG kinetics may deviate from the steady state in certain short wakeful or sleep segments, impacting the accuracy of quantified FDG metabolic changes during NREM sleep (see simulation and discussions in supplementary methods, Fig. S5). While this potential quantification bias does not alter our conclusions regarding the spatiotemporal dependence of sleep-wake hemodynamic and metabolic changes, it can be refined in future studies to achieve more precise quantification of FDG uptake, e.g., using Patlak graphical analysis (as performed in the supplementary methods, Fig. S6) and increasing the duration thresholds for stable sleep/wake segments.

In addition to elucidating the hemo-metabolic dynamics underlying transient, periodic sensory arousal during NREM sleep, our observation of temporally coordinated hemodynamic and metabolic patterns across wakefulness and light/intermediate sleep also holds important clinical implications for sleep medicine; for instance, regarding the clearance and production of metabolic waste. Recent studies have shown that fMRI-based global vascular responses are coupled to CSF dynamics^9,10^, thus a direct corollary of our observation (Fig. 2B) is that CSF dynamics also couples to metabolic dynamics across the sleep-wake cycle, with higher CSF flow when metabolism is suppressed (Fig. 6). Given that CSF dynamics may facilitate the clearance of metabolic waste during sleep^42,46^, and that sleep-wake synaptic activity and metabolic dynamics play a pivotal role in waste production, including the accumulation of toxic substances such as amyloid and tau^12,47^, our results may imply a temporal balance of the waste clearance and production processes. This simultaneous fPET-fMRI framework hence provides an exciting opportunity to further characterize and elucidate, in humans, how disrupted coordination of these processes in sleep disorders can lead to various neurological illnesses, complementary to prior studies that investigated each mechanism in isolation. Finally, from a methodological perspective, our study also holds many exciting implications for the field of dynamic multi-modal imaging involving fPET. First, unlike existing fPET-FDG studies that deliver stimuli in a temporally sparse, well-controlled manner, our results suggest that fPET-FDG can track naturalistic metabolic variations and could be applied to map metabolic processes under broad naturalistic paradigms (e.g., movie clips or speech) that are gaining popularity in cognitive neuroscience. Second, while emerging evidence has suggested that fPET-FDG can track rapid metabolic changes approaching the temporal resolution of one minute or below^23,24^, existing PET-MRI studies have primarily focused on the static, time-averaged (de)couplings of fPET and fMRI signals^15,16^, neglecting the rich information embedded in the time-resolved cross-modal dependence. Our approach of linking the dynamic evolutions of fPET metabolic measures to simultaneously acquired, instantaneous EEG and fMRI signals fully exploits the benefits of dynamic multi-modal imaging. This framework can not only help uncover the time-resolved interplay between neuronal, hemodynamic, and metabolic processes that are otherwise obscured in a time-averaged analysis, but also allow us to probe cerebral metabolism in the absence of an explicit stimulus, e.g., during endogenous arousal or sedation, when we depend on concurrent EEG signatures to infer spontaneous neuronal events.

**Figure 6:**
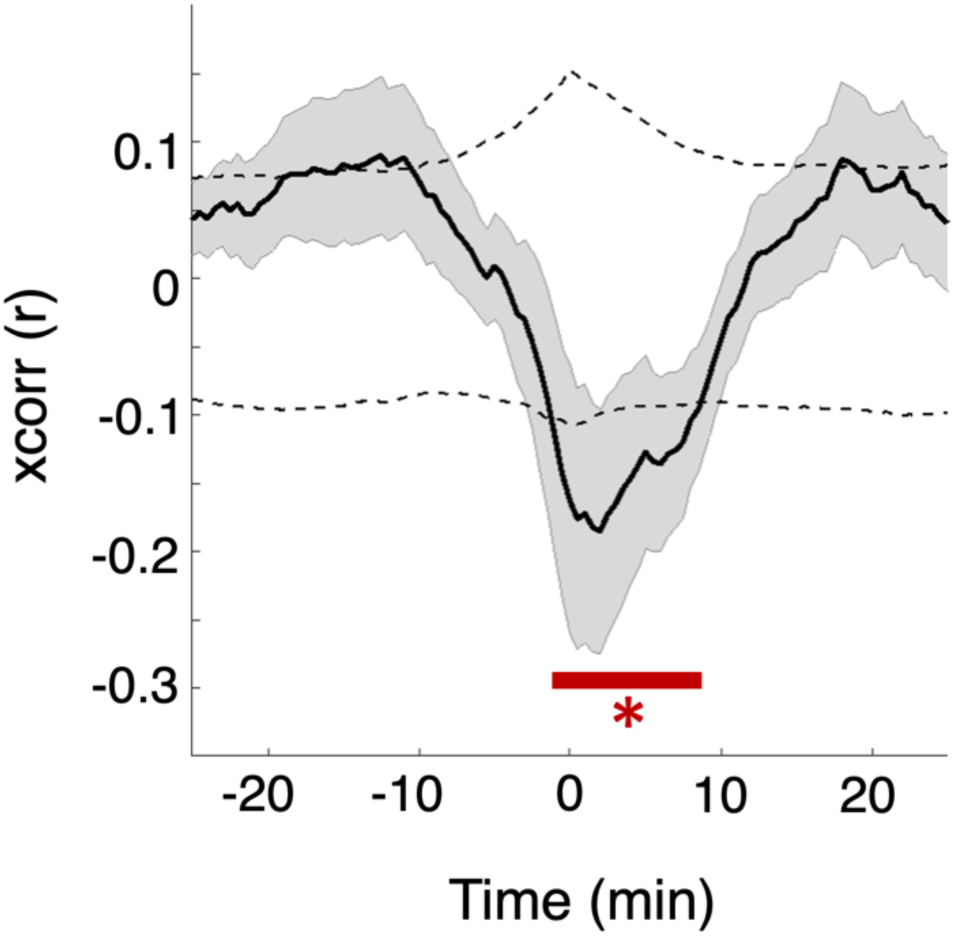
Temporal coupling between the global metabolic change and CSF dynamics across wakefulness and sleep. Cross-correlations between the global fPET-FDG signals and the metabolic TACs modeled by the amplitude variation of CSF dynamics over time, after removal of baseline trends (see ***Methods***). Positive lags indicate delayed fPET-FDG TAC relative to CSF-TAC. Despite the relatively low inflow contrast in our fMRI acquisitions due to long imaging TRs, we were able to identify a statistically significant (*p*<0.05) association between the global metabolic signal and CSF. Mean and standard error across subjects are shown (*N*=23). Gray dashed lines indicate 95% confidence intervals of the null group-averaged correlations, derived nonparametrically from phase-reshuffled data.

In summary, we show that the descent into sleep leads to coupled hemodynamic oscillations and metabolic changes across the brain. Importantly, two distinct spatial networks emerge: an oscillating, high-metabolism sensorimotor network remains active and dynamic in NREM sleep, contrasted by suppressed hemodynamic and metabolic activity in higher-order regions. These results shed light on how sleep produces a loss of awareness while preserving the brain’s ability to sensory input, and identify a complex, alternating balance of neural, vascular, and metabolic dynamics in the sleeping brain.

## Methods

### Subjects and sleep-wake experiments

Each subject participated in an (EEG-)PET-MRI session (88±11 mins), during which they were instructed to close their eyes and relax. All subjects were sleep-deprived, having only 4 hours of sleep the night before the visit to increase sleep pressure, and were asked to refrain from consuming caffeine on the day of the experiment to increase the likely range of their arousal. All subjects fasted for more than 4 hours prior to the FDG experiment. All experiments took place in the afternoon to capture the circadian afternoon dip. All studies involving human subjects were reviewed and approved by the Institutional Review Board at Massachusetts General Hospital. All subjects provided written, informed consent in accordance with the Human Research Committee at Massachusetts General Hospital. A total of 32 healthy subjects (32±8 yrs, 22 females) were enrolled in this study, among which the data of 6 subjects were excluded from the analyses due to excessive head movements (4 subjects), failure to perform the scan (1 subject), and an unexpected scan interruption due to FDG tracer leakage (1 subject). Twenty-three out of the remaining 26 subjects underwent sleep-wake transitions (i.e., including both stable wakeful and sleep conditions) during the imaging session.

### Acquisition and preprocessing

All PET-MRI experiments were performed on a 3T Tim MAGNETOM Trio MR scanner (Siemens Healthcare, Inc) with an MR-compatible BrainPET insert (Siemens). Subjects’ arousal states were delineated using simultaneous EEG recordings (21 subjects) or behavioral measures (5 subjects). For scans involving simultaneous EEG acquisitions, electrocardiogram (ECG) and respiratory traces were also collected as ancillary recordings of the EEG system throughout the sleep-wake experiments. The overall scheme of data acquisition is summarized in Fig. S1.

### MRI acquisition and preprocessing

Multi-echo MPRAGE (1 mm iso., TE1/TE2/TE3/TE4=1.64/3.5/5.36/7.22 ms, TI = 1200ms, TR = 2530 ms) images were collected to provide high-resolution T_1_-weighted anatomical image for PET photon attenuation correction and cross-modal registrations. A standard multi-slice gradient-echo EPI sequence (voxel size = 3.1×3.1×3 mm^3^, slice number = 37/44, TR = 2000/2400 ms, TE = 30 ms) was used to track brain hemodynamic changes across sleep-wake cycles. All BOLD-weighted EPI data underwent standard preprocessing steps, including corrections for motion (both rigid-body co-registration and linear regression of six translational and rotational motion parameters), slice timing, and distortion incurred by field inhomogeneities using an independent acquisition with opposite phase-encoding direction^48^.

### PET acquisition and preprocessing

Administration of FDG followed a bolus plus constant infusion paradigm (initial activity of total dose: 9.8±1.3 mCi, bolus dose was 20% of the total constant-infusion dose). FDG in saline solution was administered intravenously throughout the experiment using a Medrad Spectra Solaris syringe pump. The PET images were reconstructed with a standard 3D ordinary Poisson ordered-subset expectation maximization algorithm. Attenuation correction was performed using a pseudo-CT map derived from the high-resolution anatomical MRI data^49^. The data were reconstructed with an isotropic voxel size of 1.25-mm into a volume consisting of 153 transverse slices of 256 × 256 pixels. All volumes were smoothed using a 3D filter with a 3-mm isotropic Gaussian kernel and down-sampled to 76 slices and 128 × 128 voxels (i.e., a nominal voxel size of 2.5 × 2.5 × 2.5 mm^3^). The fPET-FDG data were binned into 30-s temporal frames. All fPET images were motion-corrected by co-registering to the middle temporal frame using the *3dvolreg* function in AFNI^50^.

### EEG acquisition and processing

For experiments involving tri-modal acquisitions (21 subjects), EEG data were recorded with MR-compatible 256-channel geodesic nets and an NA410 amplifier (Electrical Geodesics, Inc.). EEG and ancillary physiological data (respiration and ECG) were acquired at 1000 Hz and resampled to 200 Hz for preprocessing. The EEG system acquisition was synchronized with the MRI scanner’s 10 MHz master synthesizer to align data sampling with the gradient artifact, and a TTL signal was fed into the EEG system at the beginning of each volume acquisition to facilitate the denoising of gradient artifacts via template subtraction. MR gradient and ballistocardiogram artifacts were removed using the FieldTrip toolbox and a reference-layer based de-noising approach^9,51^. After removal of gradient and ballistocardiogram artifacts, EEG data were re-referenced to the mean channel for the identification of sleep stages.

Sleep staging was manually performed by an expert rater blind to study hypotheses. Each non-overlapping 30-s epoch was classified as wakeful or NREM sleep (stage N1-3) according to the AASM guideline. No REM sleep stages were identified in this study.

### EEG-based arousal index (EEG_arousal_)

Instantaneous arousal levels for each subject throughout the sleep-wake experiment were estimated using concurrent EEG measures. The power spectral density for each EEG channel was estimated using the *mtspectrumc* function from the Chronux toolbox (http://chronux.org). We calculated the ratio of the root mean square (RMS) amplitude in the 7−13 Hz range (alpha wave activity) over the RMS amplitude in the 1−7 Hz range (delta and theta activity) of signals from clean occipital EEG channels to track arousal dynamics^31^. This calculation was performed across a sequence of 30-s windows to match the temporal resolution of fPET TACs, providing a time-varying measure of arousal fluctuations over time.

### Behavioral data

For experiments not involving simultaneous EEG acquisitions (5 subjects), subjects were instructed to perform a simple breath-counting task—pressing keys on each breath in or breath out—to indicate the instantaneous arousal level. The sleep/wake state was assessed across non-overlapping 30-s epochs based on the frequency of key pressing: “wake” = continuous key pressing: “sleep” = intermittent or missing key pressing.

Experiments with either EEG or behavioral measures yielded comparable fPET-fMRI observations (see Fig. S11), and were therefore combined to provide the group-level characterizations of metabolic and hemodynamic patterns across sleep-wake cycles.

### Cross-modal registration and brain parcellations

All analyses were performed in the subject’s native space, and co-registrations between the high-resolution MR anatomical image and all functional scans (including fPET and fMRI datasets) were estimated using Freesurfer’s *bbregister* function. To enhance the contrast-to-noise ratio for cross-modal comparisons, all the analyses were performed at the region of interest (ROI) level. All brain ROIs were delineated using the high-resolution anatomical data and then re-registered to the native image space of the fPET-FDG and fMRI data. No additional spatial smoothing was performed for both datasets.

A multi-resolution functional-connectivity-based parcellation atlas was employed to delineate cortical gray-matter ROIs^52^. This multi-resolution atlas allowed us to characterize functional and metabolic changes at different spatial resolutions, and both 100-parcel and 300-parcel cortical atlases were examined in this study. Subcortical structures were segmented using the “aseg” anatomical parcellation of Freesurfer^53^. Individual white-matter bundles were obtained by renormalizing the HCP X-tract atlas^54^ to each subject’s native anatomical space using ANTs^55^. To mitigate the potential partial volume contamination from gray matter, all white-matter ROIs were further constrained by an eroded white-matter mask of the aseg atlas. After delineating all brain ROIs using the high-resolution anatomical data, the ROIs were again registered to the native image space of the fPET-FDG and fMRI data to characterize region-specific hemodynamic and metabolic patterns.

### Quantifying sleep-wake hemodynamic and metabolic changes

The amplitudes of BOLD fluctuations across sleep-wake cycles were quantified to characterize sleep-induced changes in hemodynamics. We first estimated the temporal standard deviation of fMRI signal intensities across the sequences of non-overlapping 30-s windows, yielding a dynamic time course of the amplitude variations (AVs) of BOLD fluctuations; we then computed the fractional changes of mean BOLD-AV in NREM sleep compared to wakeful states. Note that we quantified the amplitudes of hemodynamic fluctuations using the temporal standard deviations of fMRI signal intensities, rather than the peak-to-valley differences, such that the results were not dependent on the accuracy of peak/valley detections.

Next, to estimate sleep-induced changes in glucose metabolism, we quantified the fractional changes in the slopes of fPET-FDG TACs during NREM sleep compared to wakeful states. No temporal detrending was performed for this analysis. The rationale behind this analysis is that, under the steady-state condition, the slopes of the fPET TACs are proportional to the net influx rate of FDG^22^.

In order to achieve stable hemodynamic and metabolic measures in each state, all discrete-state analyses were focused on awake and NREM sleep segments which were long and stable (>5-min continuous wake or NREM sleep epochs), and data points of the initial five minutes of each state segment were excluded from the analysis.

### Inflow contrast-based sleep-wake CSF dynamics

CSF time course was extracted from the BOLD-fMRI data by averaging the signals of voxels within the fourth ventricle (segmented using the FreeSurfer “aseg” atlas), following Fultz et al. ^9^. Despite the long imaging TR of our fMRI acquisitions, we were able to detect CSF flow dynamics, evidenced by the replication of CSF-global fMRI coupling patterns reported previously^9^ (see Fig. S12). CSF-AV, the instantaneous amplitude variation of CSF fluctuations, was estimated by taking the temporal standard deviation of CSF signals across the sequences of non-overlapping 30-s windows, akin to BOLD-AV.

### Cross-modal analysis and null models

#### Linking BOLD-AV, EEG_arousal_, and CSF-AV with fPET TACs

To further link these hemodynamic, EEG arousal measures, and fluid dynamics to fPET-derived metabolic dynamics, we calculated the temporal integral of the windowed BOLD-AV, CSF-AV, and EEG_arousal_ measures, mimicking the FDG accumulation over time due to metabolic demand. Long-term baseline trends from both the modeled and measured fPET TACs were fitted and removed using a third-order polynomial, to emphasize the dynamic features of metabolic information.

After unifying the time scales of different measures, we performed temporal cross-correlation analyses to quantify cross-modal couplings and their temporal lags, and we further performed linear regression analyses to characterize the detailed spatial patterns of cross-modal dependence. To account for potential nonlinearity in the regression analysis, a square term of the regressor was also included, which however did not significantly improve the fitting results (Fig. S2).

#### Construction of null models

Null distributions of all across-modal associations (including both cross-correlation and regression analyses, global and parcel-wise analyses) were constructed nonparametrically through temporal-autocorrelation-preserved permutation tests. Briefly, for each signal pair of each individual’s data, we first generated a set of sham signal pairs by randomly reshuffling the phases of the signals in the frequency domain, we then performed the cross-correlation and regression analysis on the sham time courses. This process was repeated 5,000 times to construct the null distributions of group-level cross-modal associations.

## Author Contributions

J.E.C., B.R.R., C.Y.S., L.D.L., and C.Catana conceived and designed the study. J.E.C. and K.S.D. performed the experiment. J.E.C. and S.E.C. performed the formal analyses with support from J.F., J.C.P., C.Y.S., R.P., C.Catana, H.Y.W., J.R.P., L.D.L., and D.S.M.. L.D.L., C.Chang, and D.S.M. helped interpret the results. B.R.R., J.C.P., and C.Y.S. supervised the study. B.R.R., J.E.C., and J.R.P. funded this study. J.E.C. wrote the manuscript. L.D.L., B.R.R., C.Y.S., J.C.P., J.R.P., C.Chang., D.S.M., C.Catana and S.E.C. edited the manuscript.

## Acknowledgments

We would like to thank Shirley Hsu, Grae Arabasz, Oliver Ramsey, and Amy Kendall for their help with PET-MRI scanning support, Doug Greve, Dimitrios Mylonas, and Arun Garimella for valuable discussions regarding the present findings, Thomas Witzel and Donald Straney for setting up the EEG cable connections in the PET-MRI scanner, and Nina Fultz for her help with managing the study protocol. This work was supported in part by the NIH (grants K99/R00-NS118120, R01-MH111438, P41-EB030006, R01-MH0926638, U19-NS123717, U19-NS128613, and R21-MH135201), by the Harvard Mind Brain Behavior Faculty Research Award, by the Brain & Behavior Research Foundation Young Investigator Grant, by the BrightFocus Foundation Research Grant, and by the MGH/HST Athinoula A. Martinos Center for Biomedical Imaging; and was made possible by the resources provided by NIH Shared Instrumentation grants (S10-RR022976, S10-RR019933, S10-OD010759). Computational resources were generously provided by the Massachusetts Life Sciences Center (https://www.masslifesciences.com/).

## Supplementary Material

**Fig. S1.**
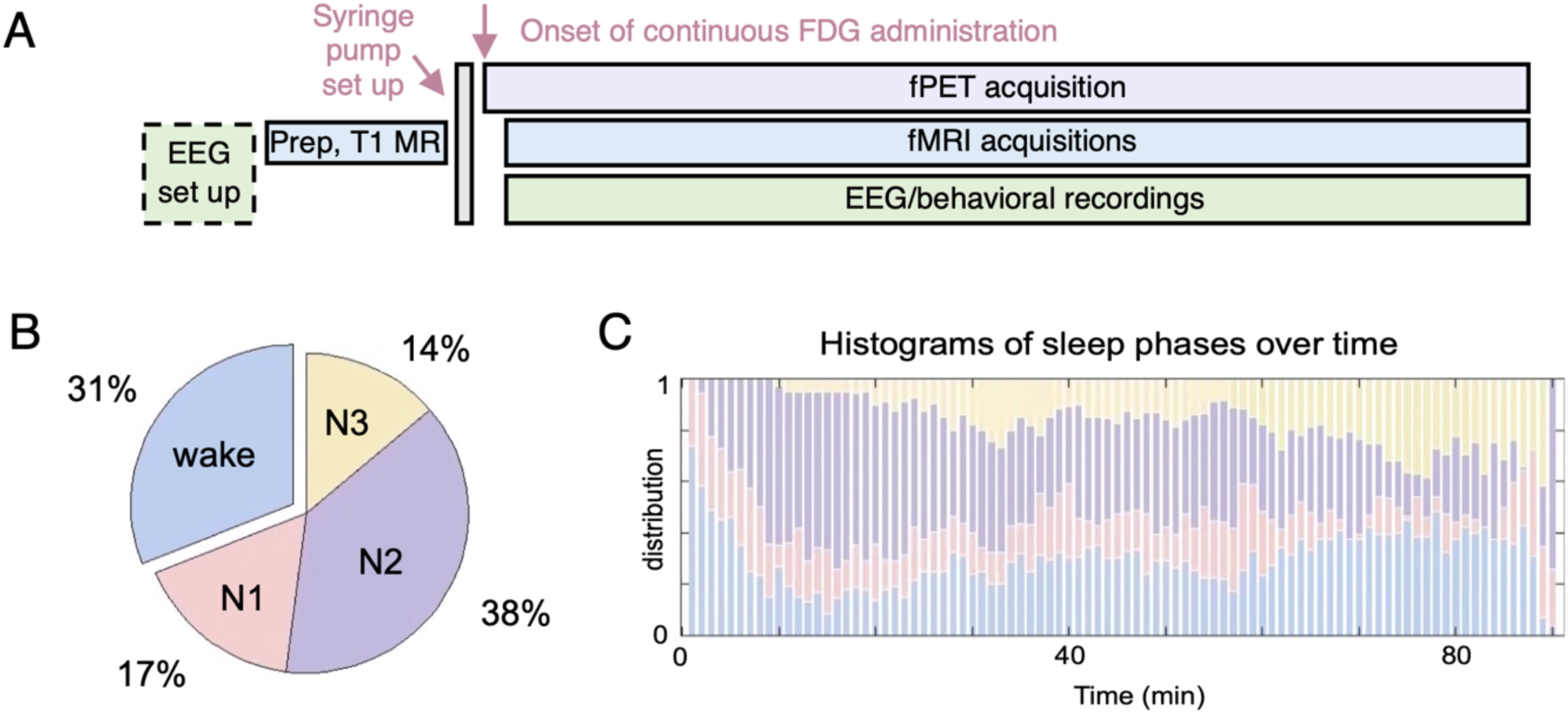
Summary of the imaging paradigms and sleep scoring statistics. (A) The overall scheme of the fPET-fMRI framework, with a total acquisition time of 88±11 mins. Among all the 26 subjects, 21 subjects underwent concurrent EEG acquisitions, and 5 subjects performed a behavioral task throughout the experiment to indicate instantaneous arousal. (B) Distributions of time spent on different arousal states, averaged across 21 subjects. (C) Histograms of sleep phase distributions over the course of the experiment, estimated across the 21 subjects with simultaneous EEG acquisitions.

**Fig. S2.**
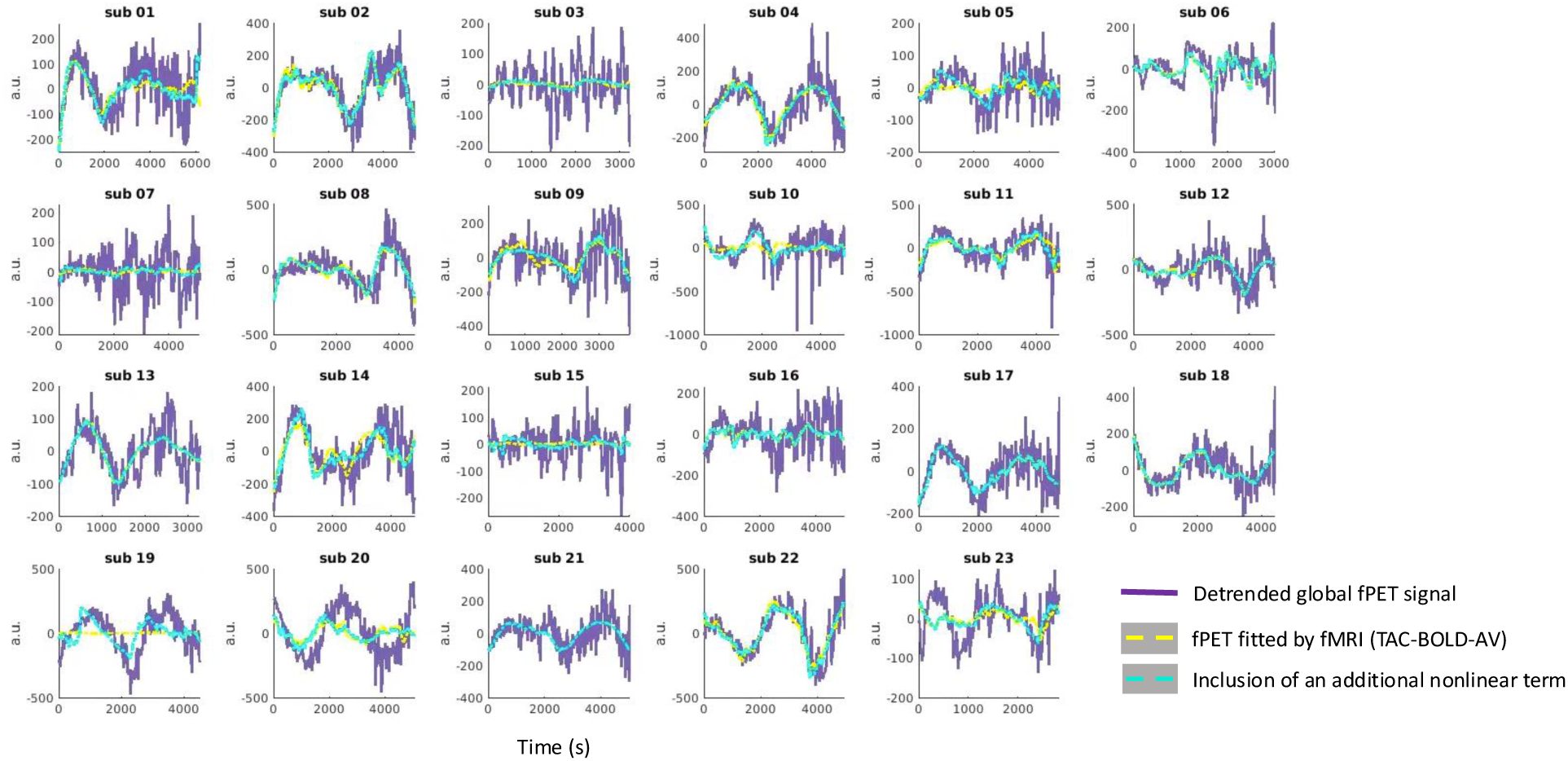
Temporal coupling between the global hemodynamic and metabolic changes at the individual level. Global fPET-FDG TACs (purple) vs. quasi-metabolic TACs modeled by the global BOLD-AV (yellow) vs. TACs modeled with the inclusion of an additional nonlinear term (cyan, the square of BOLD-AV-TACs), post temporal detrending (using third-order polynomials). Note that the inclusion of the additional higher-order non-linear term only led to modest improvements in model fitting. Results of 23 subjects undergoing sleep-wake transitions are shown.

**Fig. S3.**
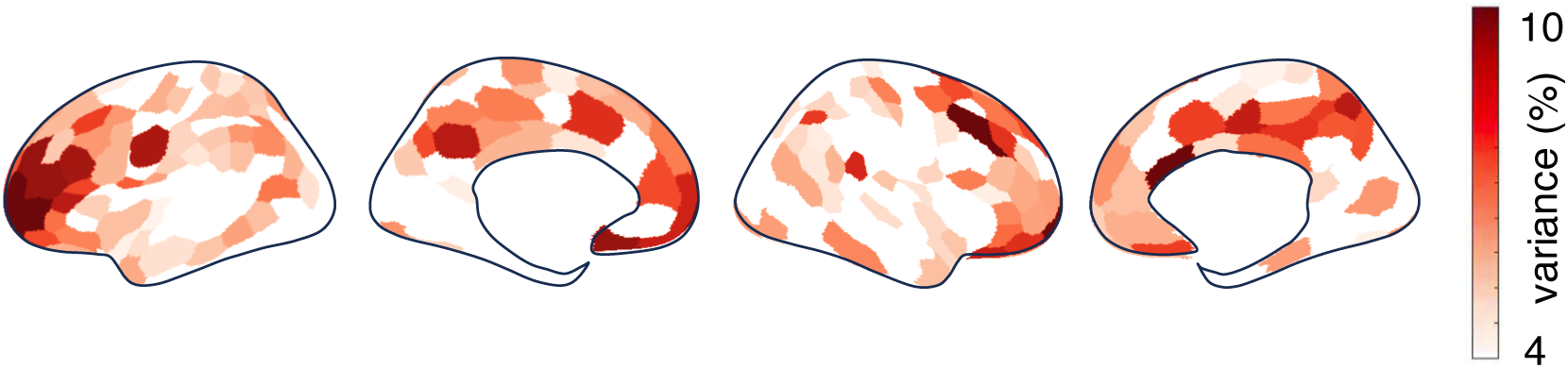
Temporal coupling between region-specific hemodynamic and metabolic changes across wake and sleep. Spatial distributions of the variance of fPET-FDG TACs explained by BOLD-AV-TAC of the same cortical parcel (*N*=23, FDR, *p*<0.05). ROI-wise fPET-FDG TACs were temporally smoothed with a median filter using a 2.5-minute (five temporal frames) kernel before variance fitting; identical temporal smoothing was applied to sham data to construct null distributions.

**Fig. S4.**
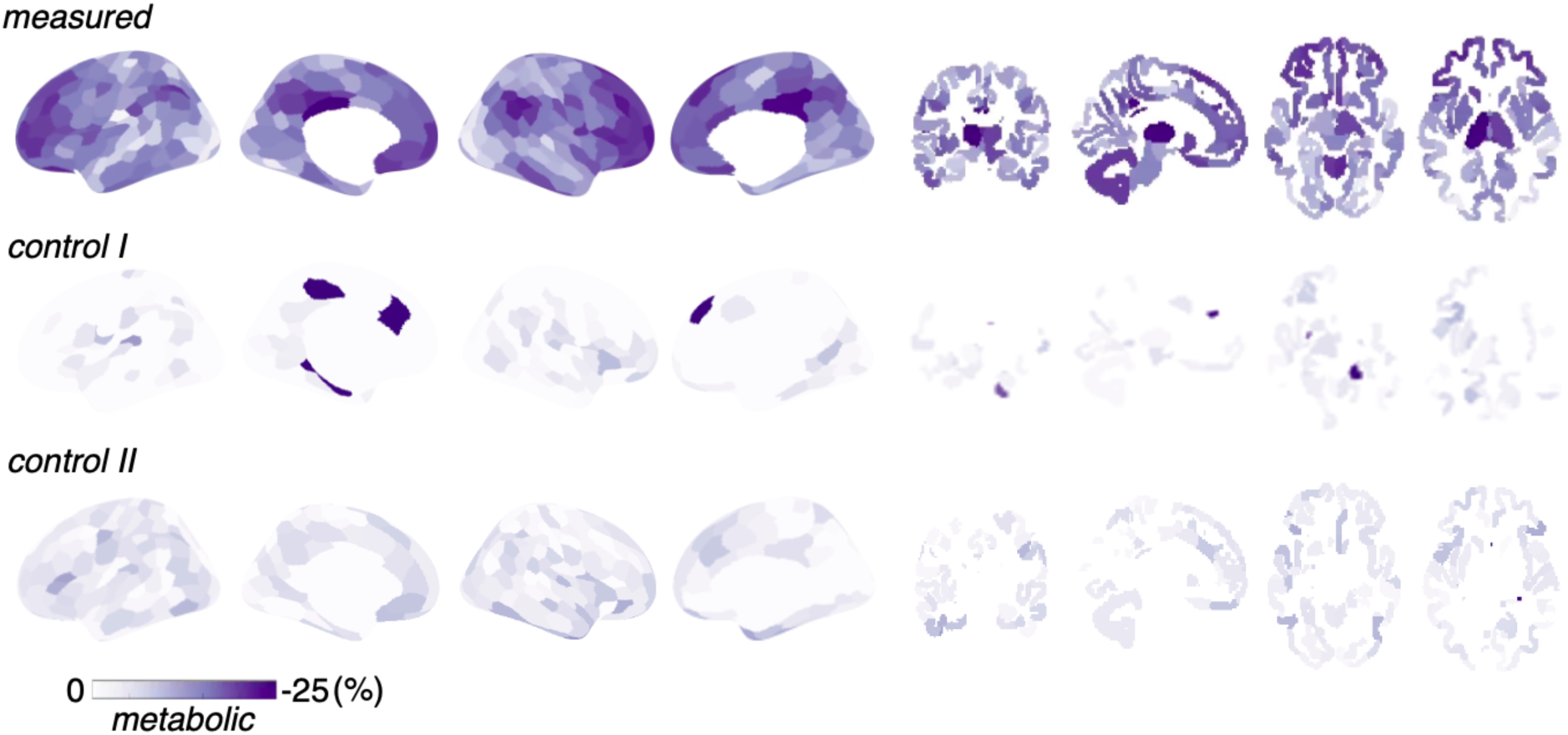
Fractional changes in FDG-based glucose metabolism during NREM sleep compared to wakeful resting: a control analysis. “Measured”: fractional changes in FDG metabolism, estimated using each individual’s fPET-FDG TACs and sleep scoring results (Fig. 3A). “Control I”: fractional changes in FDG metabolism, estimated by applying each subject’s “wake” and “sleep” scoring results to the fPET-FDG TACs of a subject that stayed awake for 96% of the time during the experiment (a “null” condition). Note that in the control analysis, since sleep-wake distributions from 23 subjects were applied to the fPET data of a single subject (eliminating inter-subject variability), the expected detection power of true effects, if exist, in the “control” case should be higher than that of the “measured” case. Additionally, while a few cortical patches exhibited apparent sleep effects, they were not statistically significant (for FDR < 0.05) and the spatial patterns altered when applying the control analysis to the fPET-FDG data of a different subject (“control II”, who drifted between Stage N1 and N2 sleep for 96% of the time during the experiment). Together, these results suggest that the measured sleep-wake metabolic changes did not stem from baseline misspecification, i.e., potential bias introduced by the interaction of sleep-wake timing and the shapes of baseline fPET-FDG TACs canceled out at the group level.

### Supplementary Methods: quantifying the fractional changes in glucose metabolism during NREM sleep compared to wakeful resting using blocked Patlak Plot

Patlak graphical analysis is a well-established method for quantifying glucose metabolism in FDG-PET studies^1^. This method, based on the compartmental models of tracer kinetics, uses a Patlak plot to determine the metabolic rate of FDG during periods of metabolic equilibrium, with the slope reflecting the net influx rate of FDG. Blocked Patlak analysis here refers to the method wherein a Patlak plot is coded into sections across which slopes are fit simultaneously and then compared. Analogous approaches have been proposed and validated previously^2,3^. Here, we applied the blocked Patlak analysis to quantify the ratio of FDG metabolism between sleep and wakefulness, which can be equated to the ratio of Patlak slopes during these respective states. Plasma arterial input functions were derived using an image-derived approach, based on the segmentation of carotid arteries from MPRAGE images^4^.

We first conducted noise-free simulations to assess potential biases in quantifying metabolic changes using the simplified approach that directly estimates the ratio of fPET TAC slopes between states (“TAC slope” method, see ***Methods*** in the main text) and using the blocked Patlak plot. An irreversible two-tissue compartmental model with bolus-plus-constant-infusion input was employed to simulate fPET-FDG TACs, with kinetic rate constants (*K_1_* = 0.1 mL/min*, k_2_* = 0.15 min^−1^*, k_3_* =0.08 min^−1^, *k_4_* = 0 min^−1^) taken from previous literature^5^. Image-derived plasma arterial input functions and sleep-wake state distributions (including the > 5-minute stable sleep/wake segments) were extracted from each subject’s experimental data. Sleep-induced fractional changes in FDG metabolism were assumed consistent across subjects and simulated by varying the *k_3_* parameter (fractional changes from -50% to 0% to simulate metabolic reductions in sleep). Overall, the Patlak analysis provided more precise estimates of FDG metabolic changes than the TAC slope approach (Fig. S5), suggesting that a longer equilibrium time is necessary to approximate the steady state for the TAC slope method (where the plasma FDG concentration becomes constant and the fPET TAC is quasi-linear) compared to the Patlak graphical analysis (where the plot converges to a straight line).

Figure S6A illustrates the blocked Patlak analysis methodology applied to real data, exemplified using two subjects’ mean cerebral gray matter TACs. Figure S6B displays the group-level results of using Patlak plots to analyze the fractional change in FDG metabolic rate between wakefulness and NREM sleep, which exhibited a general concordance with the spatial distributions of metabolic changes characterized by the TAC slope method in Fig. 3A (Spearman’s spatial correlation = 0.73, *p* < 1e-15, 300 ROIs). This suggests that, while the TAC slope-based method may not yield precise quantification, it is able to estimate the qualitative spatial distributions of FDG metabolic changes in our study, owing to the large effect size of sleep modulation.

**Figure S5.**
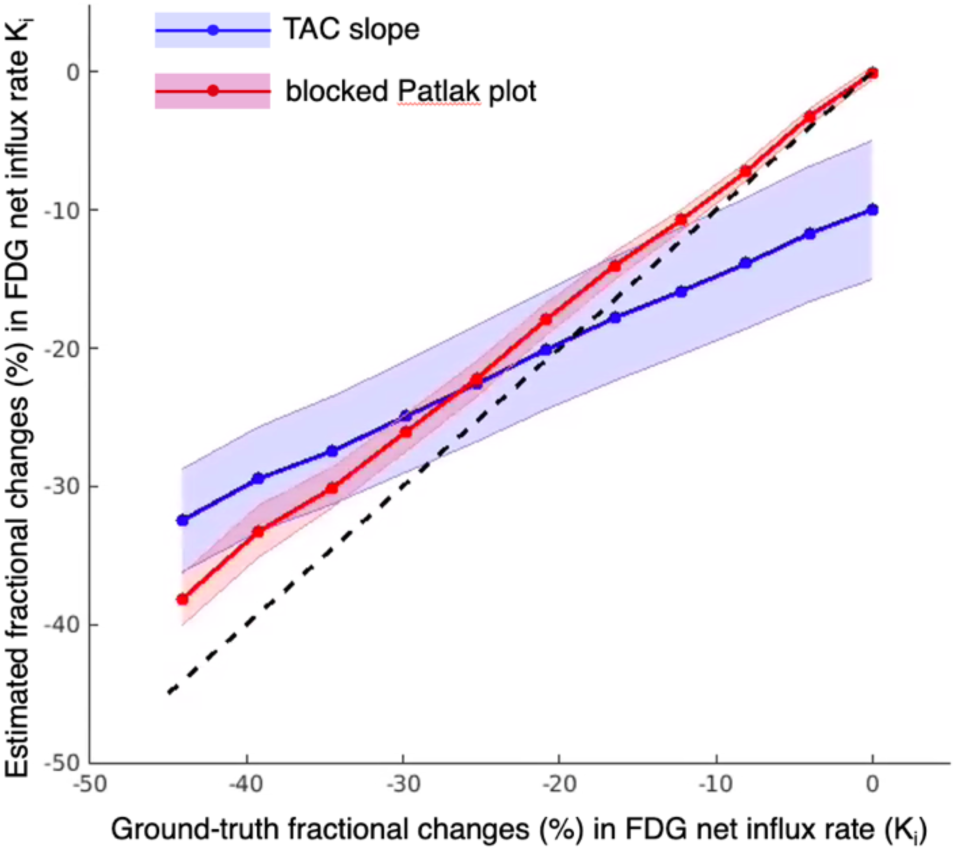
Simulation results: NREM sleep-induced fractional changes in FDG uptake. Groundtruth changes were simulated by varying the *k_3_* parameter, and the FDG net influx rate *k_i_* = *k*_1_*k*_3_/(*k*_2_ + *k*_3_). Estimated FDG metabolic changes were derived from both the simplified fPET TAC slope method (“TAC slope”) and the blocked Patlak graphical analysis (“blocked Patlak plot”); mean and standard error of estimates based on each subject’s wake/sleep distribution are shown.

**Figure S6.**
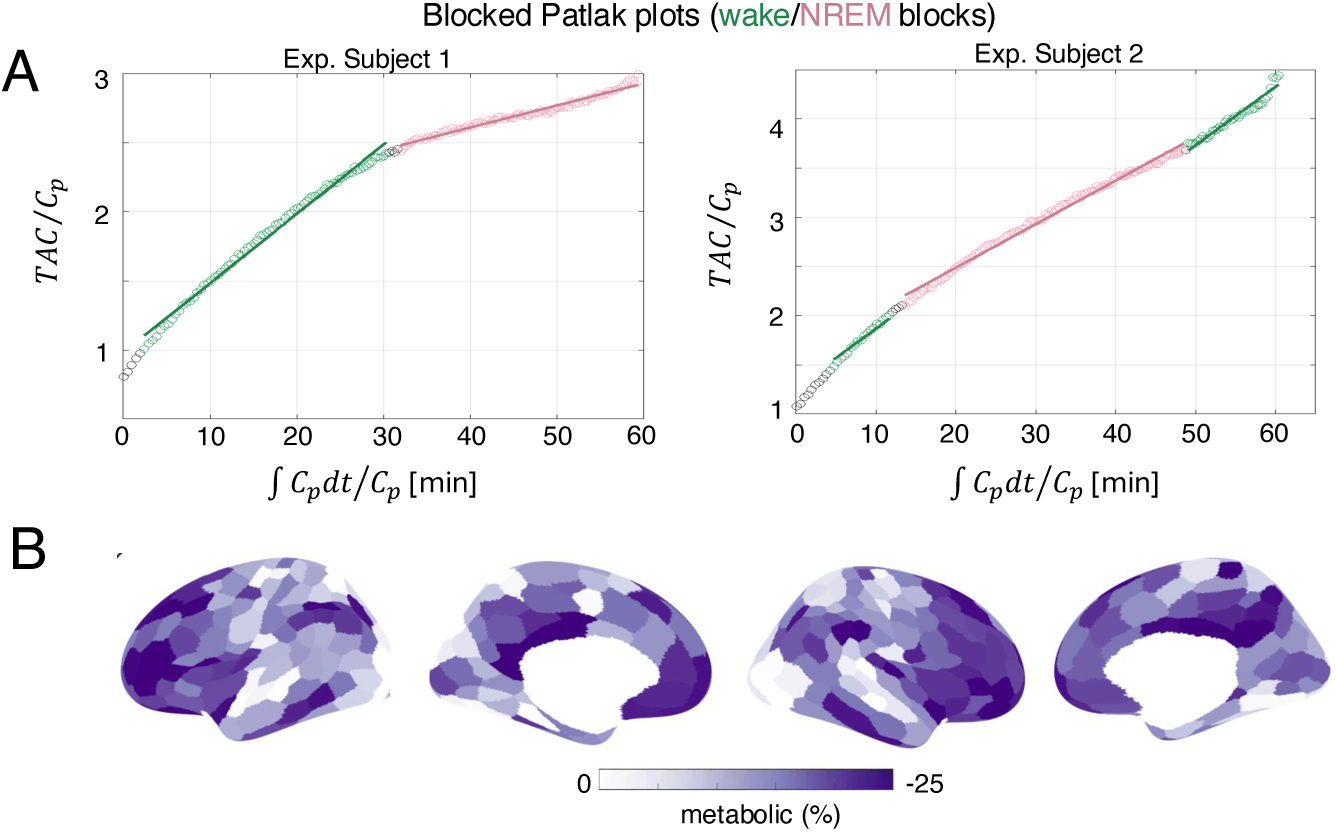
Real-data results: NREM sleep-induced fractional changes in FDG uptake, estimated using the Patlak graphical analysis. (A) Illustrative fitting of blocked Patlak plot, global mean TACs in two exemplar subjects. C_p_ indicates plasma FDG concentration. (B) Fractional change in FDG metabolism during sleep as calculated using Patlak graphical analysis, weighted average across subjects by inverse of percent error of individual estimates.

**Fig. S7.**
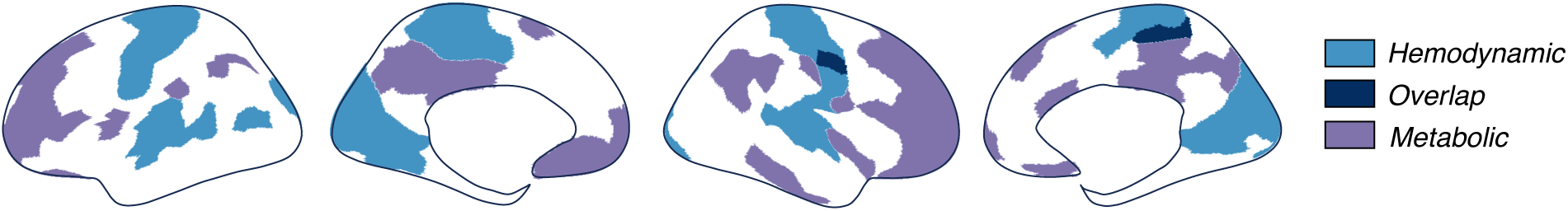
NREM sleep-induced changes in hemodynamic and metabolic signals, overlaid on the same surface. NREM sleep-induced fractional increases in BOLD-AV (blue) and decreases in FDG metabolism (purple). Cortical ROIs demonstrating the top 25% absolute changes for each modality are shown; and these cortices with the strongest sleep effects exhibit very modest overlap (navy).

**Fig. S8.**
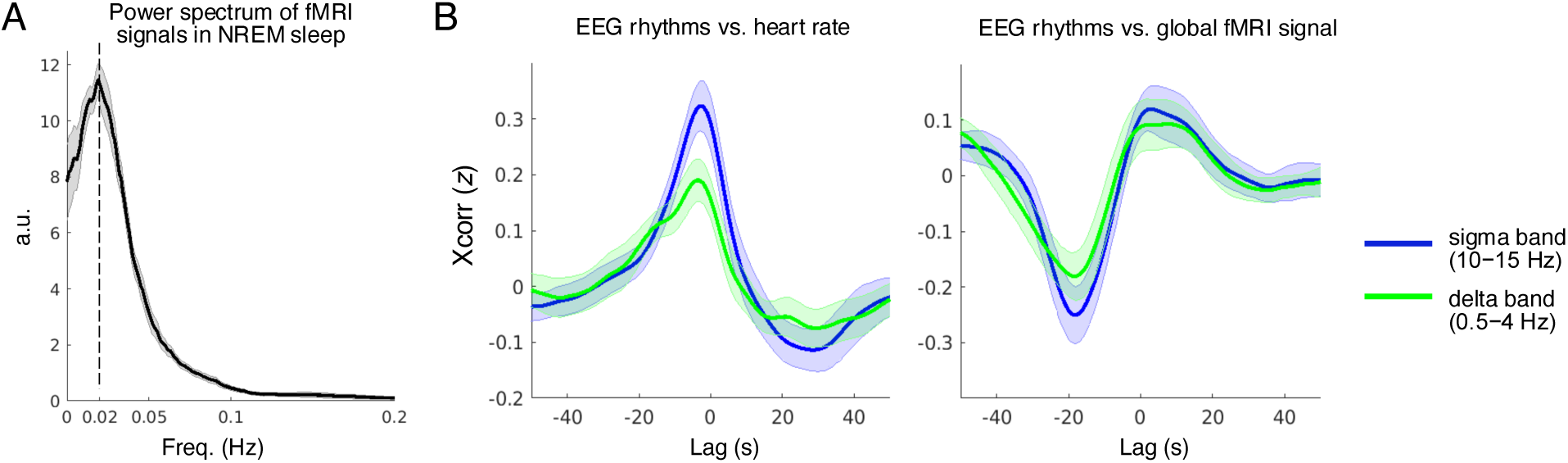
Temporally coupled large fMRI oscillations, EEG sigma band activity, and heart rate in NREM sleep. (A) FMRI oscillations in NREM sleep peaked at ∼0.02 Hz. (B) Cross-correlations between the RMS amplitude of EEG sigma-band activity (10−15 Hz) and heart rate, and the global fMRI signal; correlation results for EEG delta-band slow-wave activity (0.5−4 Hz) are also included for reference. EEG rhythms precede heart rate and global fMRI oscillations. Mean and standard errors across subjects (*N*=17). These observations align with findings from previous NREM sleep studies: (i) coordinated EEG/ECoG sigma-band activity and heart rate at 0.02 Hz in rodents and humans^6^; (ii) coupled fMRI oscillation and spindle activity in humans^7^; and (iii) coupled fMRI oscillation, slow-wave activity, and heart rate in humans^8^.

**Fig. S9.**
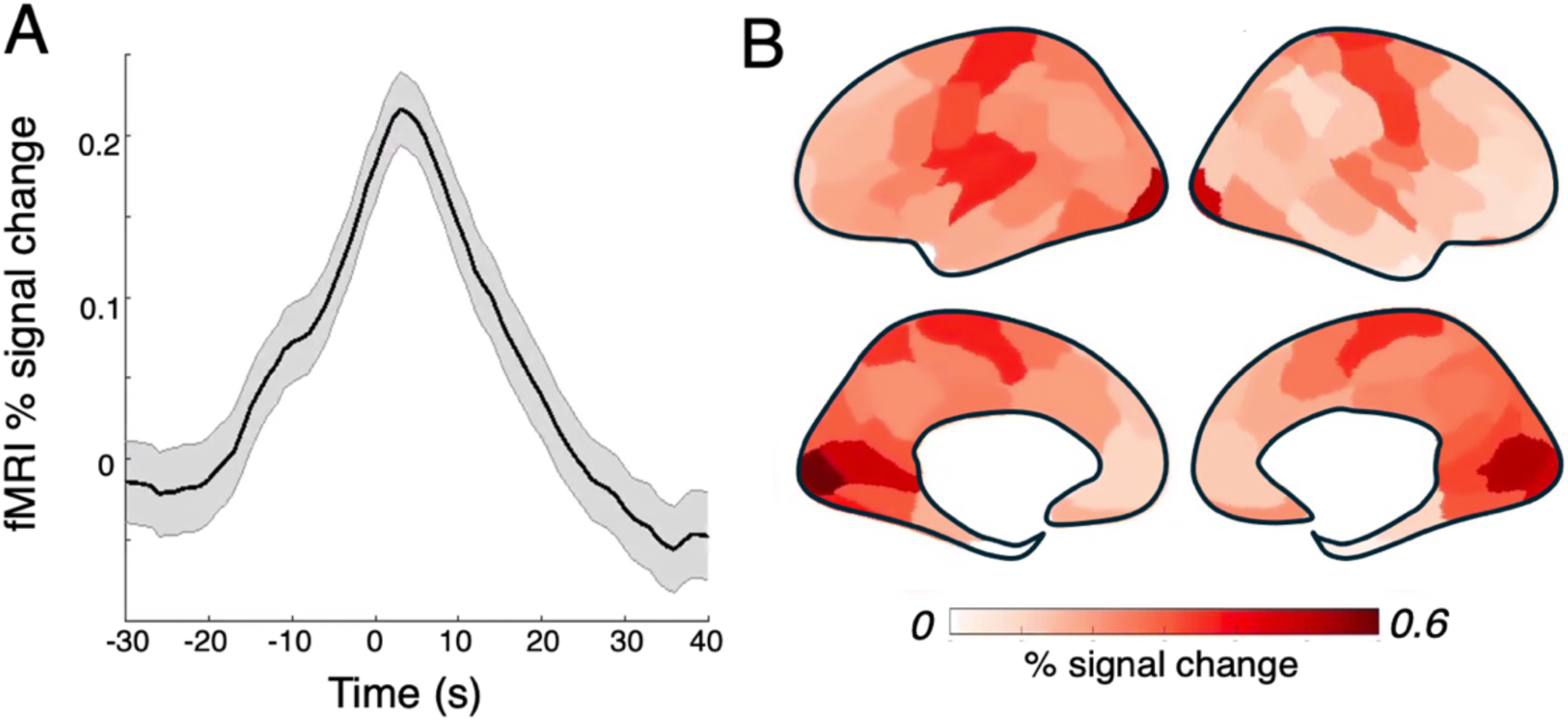
FMRI responses time-locked to discrete spindles in NREM sleep. (A) % change in global fMRI signal following a sleep spindle (time 0: center of spindle; mean and standard errors across 663 spindle events); (B) Spatial distribution of peak fMRI % signal change following a sleep spindle; note that the sensory network exhibited the largest fMRI signal change.

**Fig. S10.**
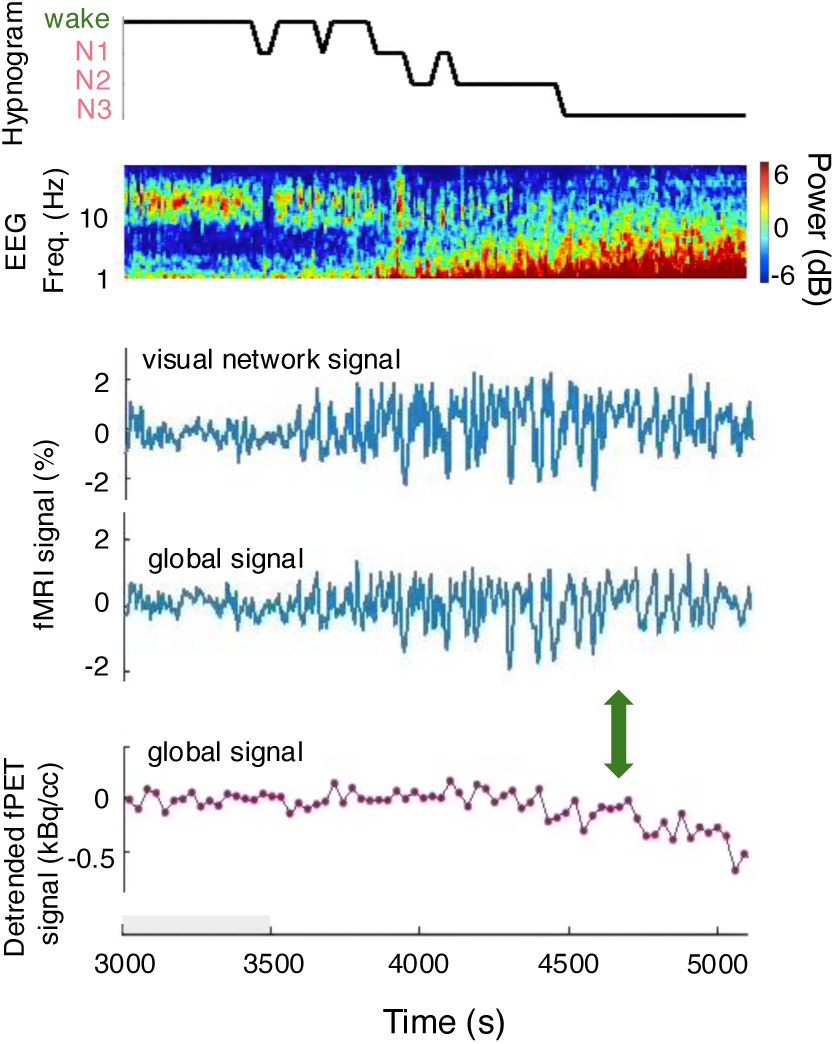
An illustrative example that both the amplitude of hemodynamic oscillations and glucose metabolism decreased as the subject gradually descended from Stage N2 to N3 sleep. Top: hypnogram of scored sleep staging and the spectrogram of an occipital EEG electrode; middle: fMRI-based global hemodynamic oscillations, the signal of the visual network is also displayed as reference; bottom: fPET-based global metabolic dynamics. Functional PET signals were detrended according to the initial wakeful period (shaded gray area) to help visualize altered slopes at state transitions. Note that the decrease in fPET TAC slope in Stage N3 sleep was not accompanied by an increase in the amplitude of fMRI oscillations (highlighted by the green arrow).

**Fig. S11.**
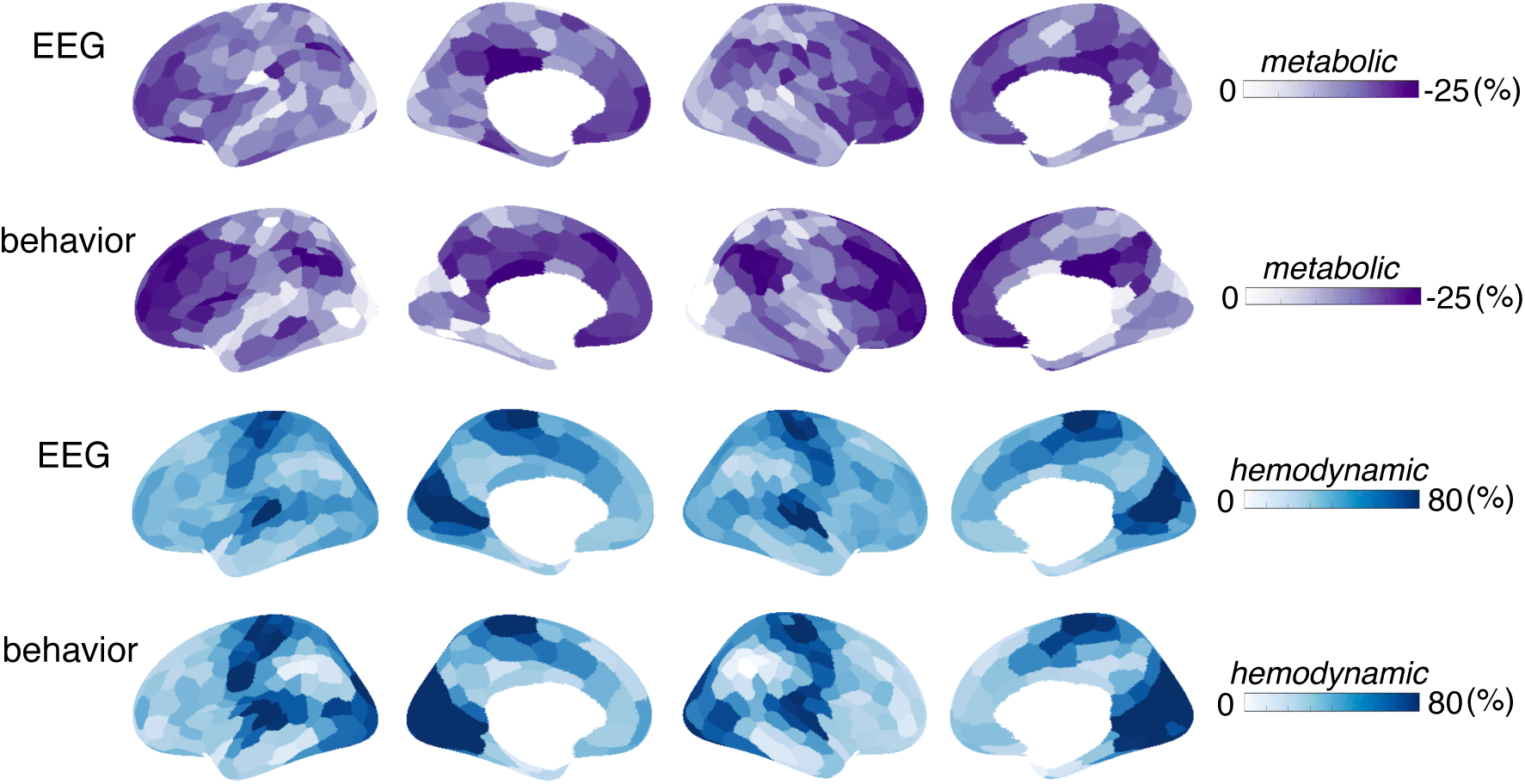
Spatial distributions of NREM sleep-induced changes in hemodynamics and FDG metabolism: EEG (*N*=18) vs. behavioral arousal (*N*=5). Overall, the EEG-based results are consistent with the main text (Fig. 3A), given the dominant number of subjects. Results yielded from EEG and behavioral arousal measures exhibit consistent spatial distributions (Spearman’s spatial correlation is 0.97 for fMRI-based hemodynamic changes; and 0.87 for fPET-based metabolic changes): sensory regions exhibit the strongest BOLD-AV changes in NREM sleep compared to wakeful states; whereas the frontal and posterior regions exhibit the strongest changes in glucose metabolism during sleep compared to wakeful states. Therefore, considering the challenges of data acquisition, results from EEG and behavioral arousal measures were combined in this study.

**Fig. S12.**
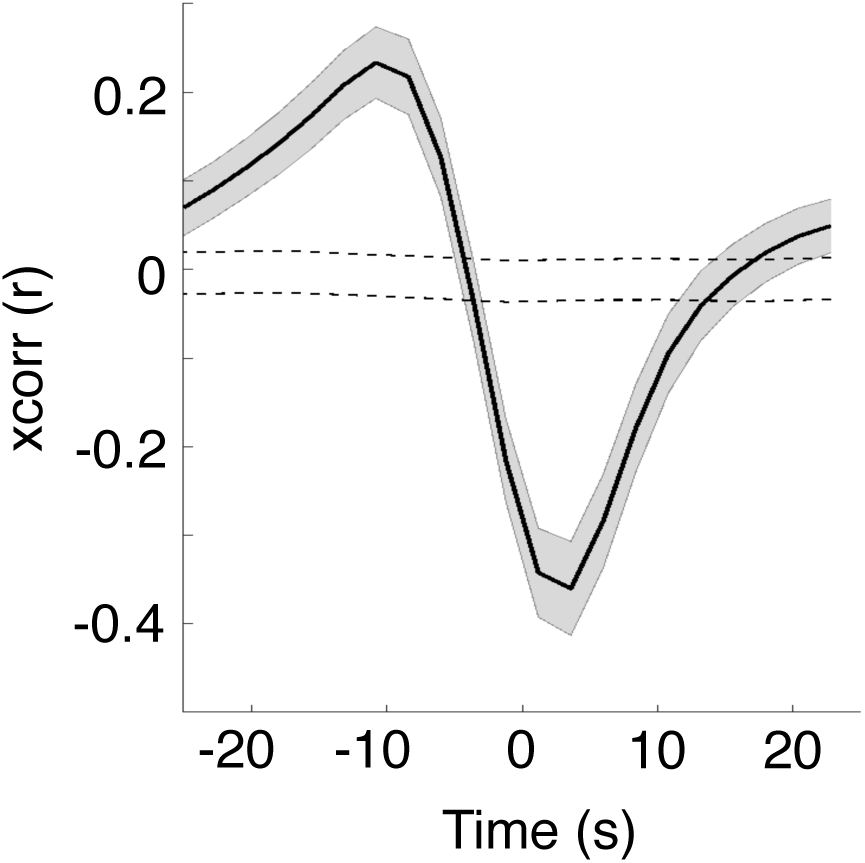
Temporal coupling between CSF and global fMRI signals. CSF time course was extracted from the BOLD-fMRI data by averaging the signals of voxels within the fourth ventricle (segmented using the FreeSurfer “aseg” atlas). Consistent with previous reports^9^, we observed a strong anti-correlation between CSF and the global fMRI signal, supporting a dominant inflow contrast despite the low imaging TR employed in the study. Positive lags indicate delayed CSF than BOLD dynamics. Mean and standard errors across subjects are shown (*N*=23). Gray dashed lines indicate 95% intervals of the null group-averaged correlations, derived nonparametrically from phase-reshuffled data.

## Notes

### Competing Interest Statement

The authors have declared no competing interest.

