## Supplementary Material for "Simultaneous EEG-PET-MRI identifies temporally coupled, spatially structured hemodynamic and metabolic dynamics across wakefulness and NREM sleep"

**Fig. S1 Summary of the imaging paradigms and sleep scoring statistics.**

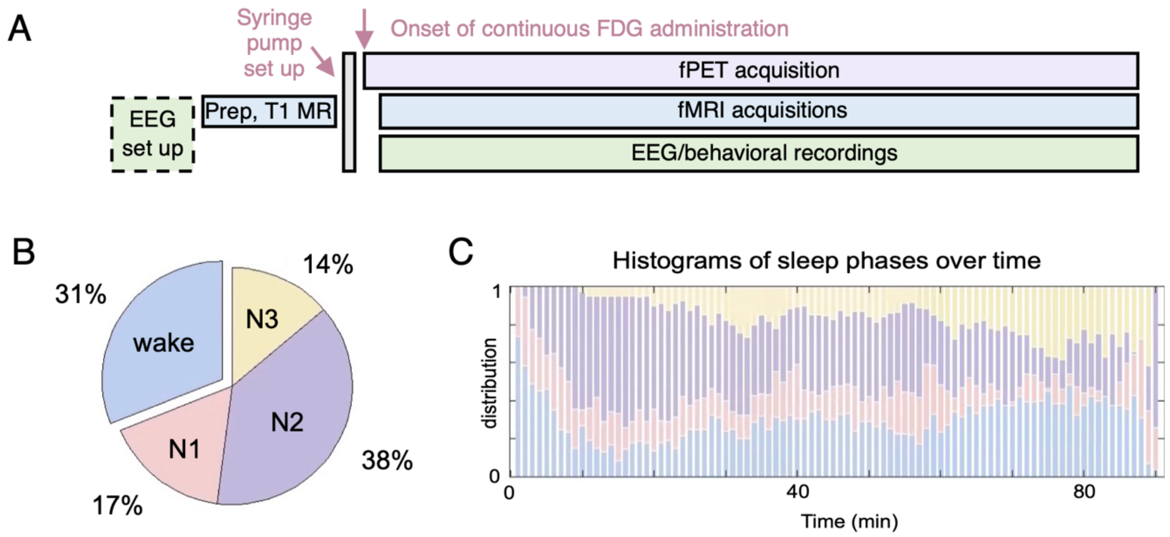


(A) The overall scheme of the fPET-fMRI framework, with a total acquisition time of 88$\pm$11 mins. Among all the 26 subjects, 21 subjects underwent concurrent EEG acquisitions, and 5 subjects performed a behavioral task throughout the experiment to indicate instantaneous arousal. (B) Distributions of time spent on different arousal states, averaged across 21 subjects. (C) Histograms of sleep phase distributions over the course of the experiment, estimated across the 21 subjects with simultaneous EEG acquisitions.

**Fig. S2 Temporal coupling between the global hemodynamic and metabolic changes at the individual level.**

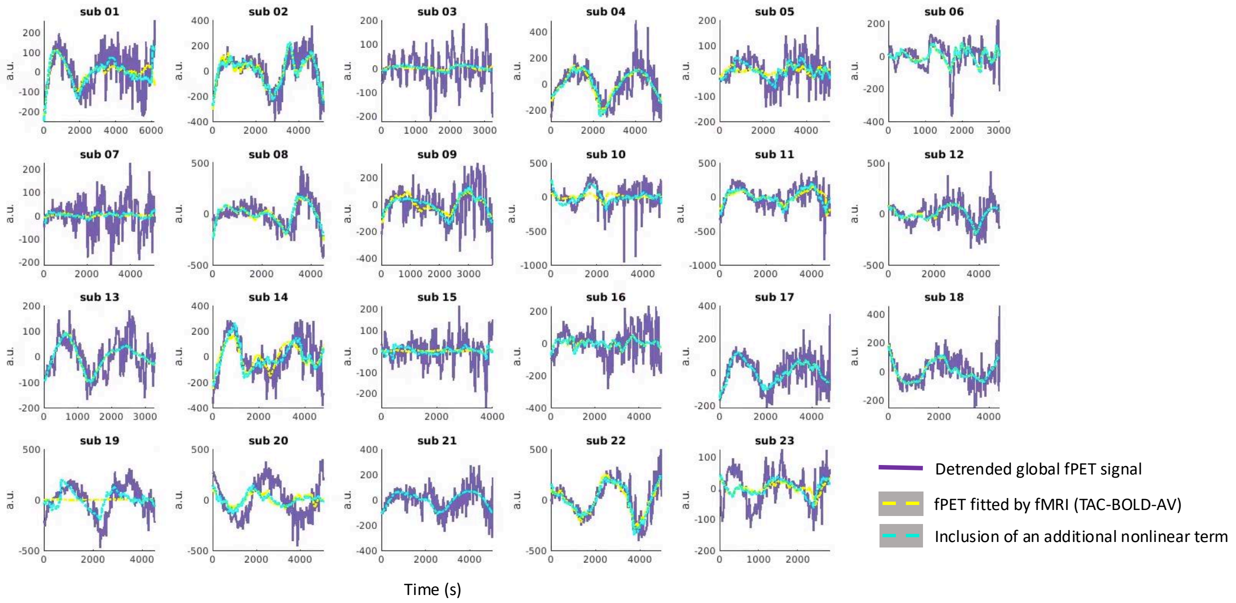


Global fPET-FDG TACs (purple) vs. quasi-metabolic TACs modeled by the global BOLD-AV (yellow) vs. TACs modeled with the inclusion of an additional nonlinear term (cyan, the square of BOLD-AV-TACs), post temporal detrending (using third-order polynomials). Note that the inclusion of the additional higher-order non-linear term only led to modest improvements in model fitting. Results of 23 subjects undergoing sleep-wake transitions are shown.

**Fig. S3 Temporal coupling between region-specific hemodynamic and metabolic changes across wake and sleep.**

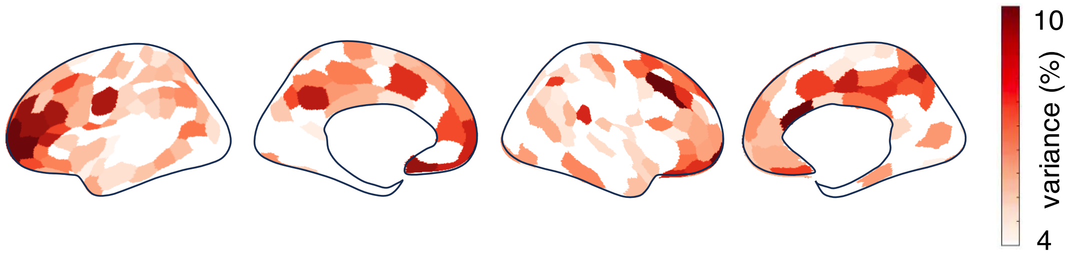


Spatial distributions of the variance of fPET-FDG TACs explained by BOLD-AV-TAC of the same cortical parcel (*N*=23, FDR, *p*<0.05). ROI-wise fPET-FDG TACs were temporally smoothed with a median filter using a 2.5-minute (five temporal frames) kernel before variance fitting; identical temporal smoothing was applied to sham data to construct null distributions.

**Fig. S4 Fractional changes in FDG-based glucose metabolism during NREM sleep compared to wakeful resting: a control analysis.**

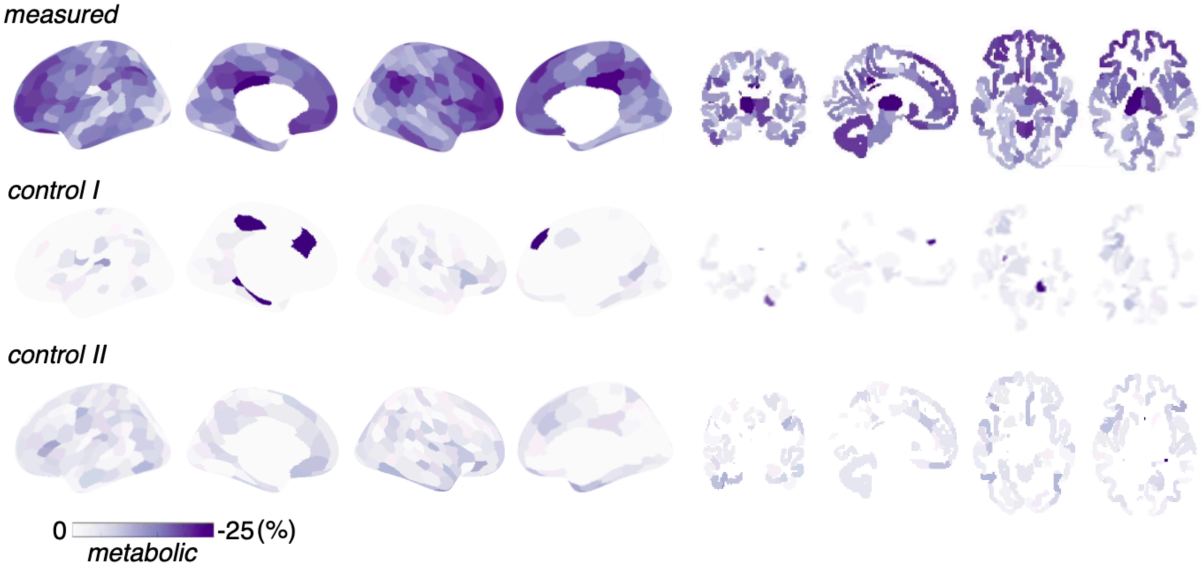


"Measured": fractional changes in FDG metabolism, estimated using each individual's fPET-FDG TACs and sleep scoring results (Fig. 3A). "Control I": fractional changes in FDG metabolism, estimated by applying each subject's "wake" and "sleep" scoring results to the fPET-FDG TACs of a subject that stayed awake for 96% of the time during the experiment (a "null" condition). Note that in the control analysis, since sleep-wake distributions from 23 subjects were applied to the fPET data of a single subject (eliminating inter-subject variability), the expected detection power of true effects, if exist, in the “control” case should be higher than that of the “measured” case. Additionally, while a few cortical patches exhibited apparent sleep effects, they were not statistically significant (for FDR < 0.05) and the spatial patterns altered when applying the control analysis to the fPET-FDG data of a different subject ("control II”, who drifted between Stage N1 and N2 sleep for 96% of the time during the experiment). Together, these results suggest that the measured sleep-wake metabolic changes did not stem from baseline misspecification, i.e., potential bias introduced by the interaction of sleep-wake timing and the shapes of baseline fPET-FDG TACs canceled out at the group level.

Figure S6A illustrates the blocked Patlak analysis methodology applied to real data, exemplified using two subjects’ mean cerebral gray matter TACs. Figure S6B displays the group-level results of using Patlak plots to analyze the fractional change in FDG metabolic rate between wakefulness and NREM sleep, which exhibited a general concordance with the spatial distributions of metabolic changes characterized by the TAC slope method in Fig. 3A (Spearman's spatial correlation = 0.73, $p$ < 1e-15, 300 ROIs). This suggests that, while the TAC slope-based method may not yield precise quantification, it is able to estimate the qualitative spatial distributions of FDG metabolic changes in our study, owing to the large effect size of sleep modulation.

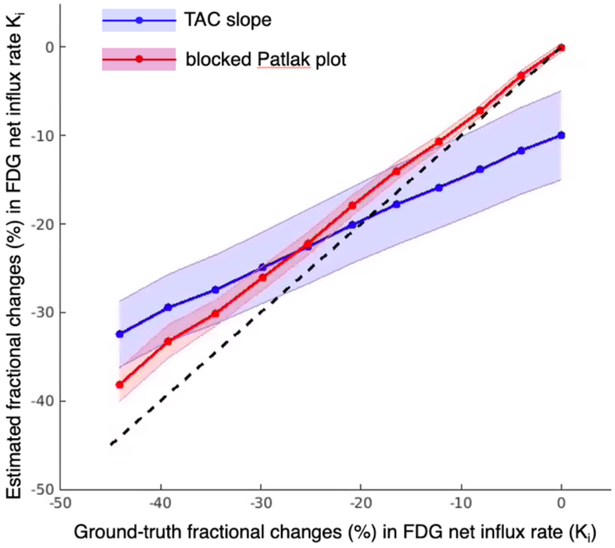


**Figure S5** Simulation results: NREM sleep-induced fractional changes in FDG uptake. Groundtruth changes were simulated by varying the $k_3$ parameter, and the FDG net influx rate $K_i = K_1 k_3/(k_2 + k_3)$. Estimated FDG metabolic changes were derived from both the simplified fPET TAC slope method ("TAC slope") and the blocked Patlak graphical analysis ("blocked Patlak plot"); mean and standard error of estimates based on each subject's wake/sleep distribution are shown.

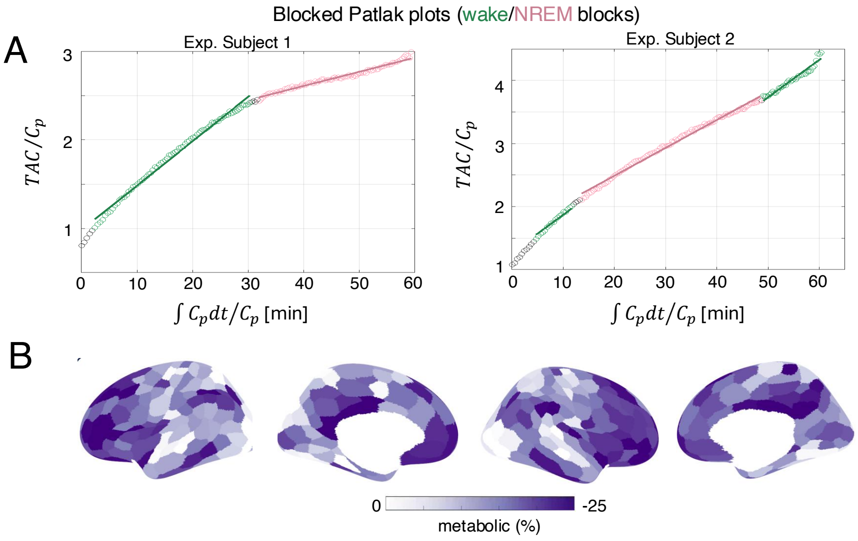

**Figure S6** Real-data results: NREM sleep-induced fractional changes in FDG uptake, estimated using the Patlak graphical analysis. (A) Illustrative fitting of blocked Patlak plot, global mean TACs in two exemplar subjects. $C_p$ indicates plasma FDG concentration. (B) Fractional change in FDG metabolism during sleep as calculated using Patlak graphical analysis, weighted average across subjects by inverse of percent error of individual estimates.

**Fig. S7 NREM sleep-induced changes in hemodynamic and metabolic signals, overlaid on the same surface.**

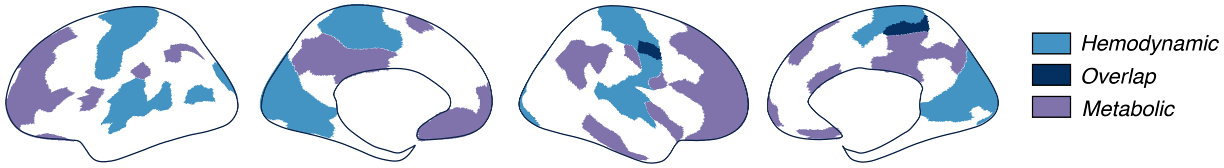


NREM sleep-induced fractional increases in BOLD-AV (blue) and decreases in FDG metabolism (purple). Cortical ROIs demonstrating the top 25% absolute changes for each modality are shown; and these cortices with the strongest sleep effects exhibit very modest overlap (navy).

**Fig. S8 Temporally coupled large fMRI oscillations, EEG sigma band activity, and heart rate in NREM sleep.**

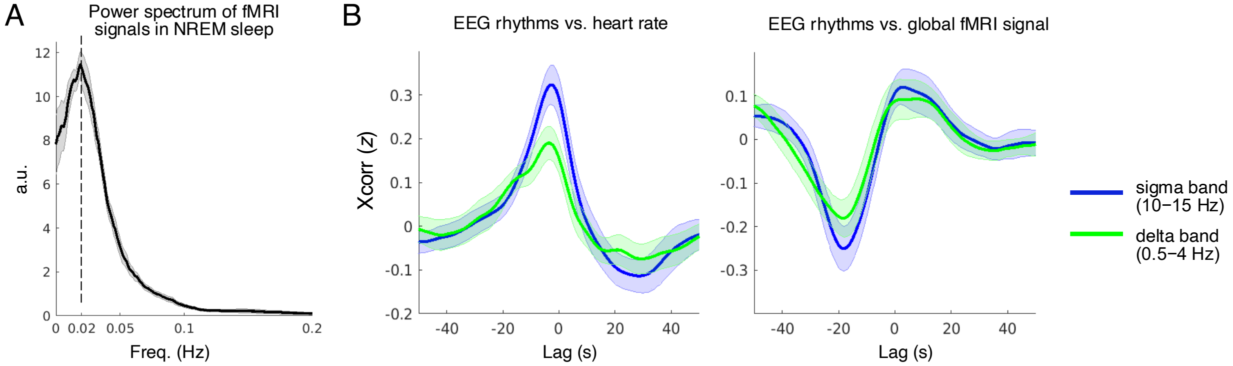


(A) FMRI oscillations in NREM sleep peaked at ~0.02 Hz. (B) Cross-correlations between the RMS amplitude of EEG sigma-band activity (10–15 Hz) and heart rate, and the global fMRI signal; correlation results for EEG delta-band slow-wave activity (0.5–4 Hz) are also included for reference. EEG rhythms precede heart rate and global fMRI oscillations. Mean and standard errors across subjects (*N*=17). These observations align with findings from previous NREM sleep studies: (i) coordinated EEG/ECoG sigma-band activity and heart rate at 0.02 Hz in rodents and humans[6]; (ii) coupled fMRI oscillation and spindle activity in humans[7]; and (iii) coupled fMRI oscillation, slow-wave activity, and heart rate in humans[8].

**Fig. S9 FMRI responses time-locked to discrete spindles in NREM sleep.**

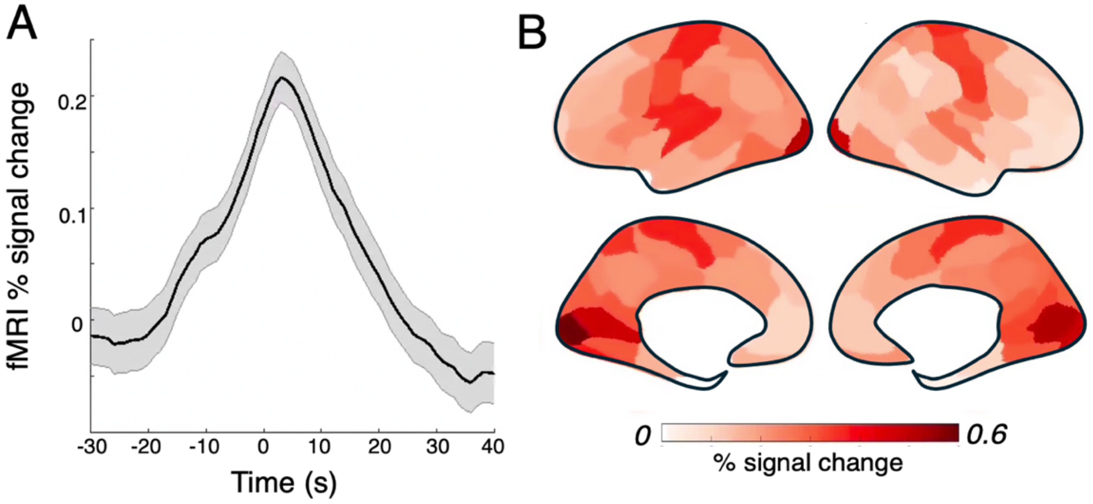


(A) % change in global fMRI signal following a sleep spindle (time 0: center of spindle; mean and standard errors across 663 spindle events); (B) Spatial distribution of peak fMRI % signal change following a sleep spindle; note that the sensory network exhibited the largest fMRI signal change.

**Fig. S10 An illustrative example that both the amplitude of hemodynamic oscillations and glucose metabolism decreased as the subject gradually descended from Stage N2 to N3 sleep.**

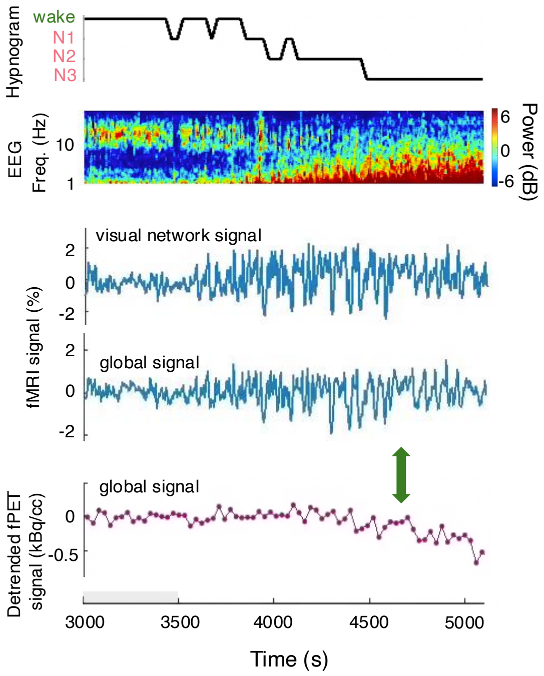


Top: hypnogram of scored sleep staging and the spectrogram of an occipital EEG electrode; middle: fMRI-based global hemodynamic oscillations, the signal of the visual network is also displayed as reference; bottom: fPET-based global metabolic dynamics. Functional PET signals were detrended according to the initial wakeful period (shaded gray area) to help visualize altered slopes at state transitions. Note that the decrease in fPET TAC slope in Stage N3 sleep was not accompanied by an increase in the amplitude of fMRI oscillations (highlighted by the green arrow).

**Fig. S11 Spatial distributions of NREM sleep-induced changes in hemodynamics and FDG metabolism: EEG (*N*=18) vs. behavioral arousal (*N*=5).**

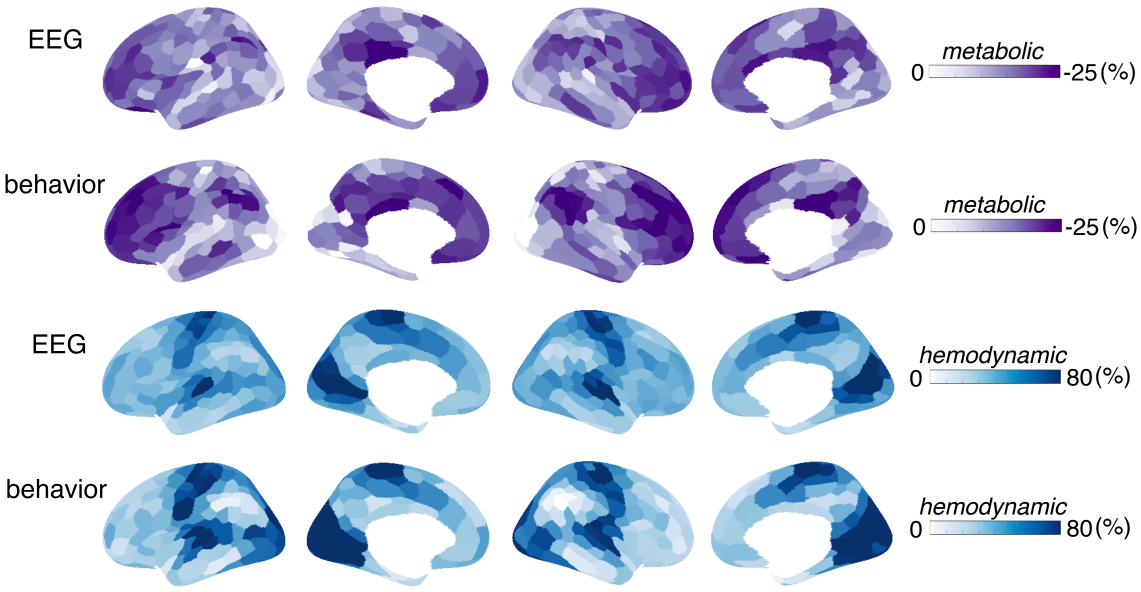


Overall, the EEG-based results are consistent with the main text (Fig. 3A), given the dominant number of subjects. Results yielded from EEG and behavioral arousal measures exhibit consistent spatial distributions (Spearman's spatial correlation is 0.97 for fMRI-based hemodynamic changes; and 0.87 for fPET-based metabolic changes): sensory regions exhibit the strongest BOLD-AV changes in NREM sleep compared to wakeful states; whereas the frontal and posterior regions exhibit the strongest changes in glucose metabolism during sleep compared to wakeful states. Therefore, considering the challenges of data acquisition, results from EEG and behavioral arousal measures were combined in this study.

**Fig. S12 Temporal coupling between CSF and global fMRI signals.**

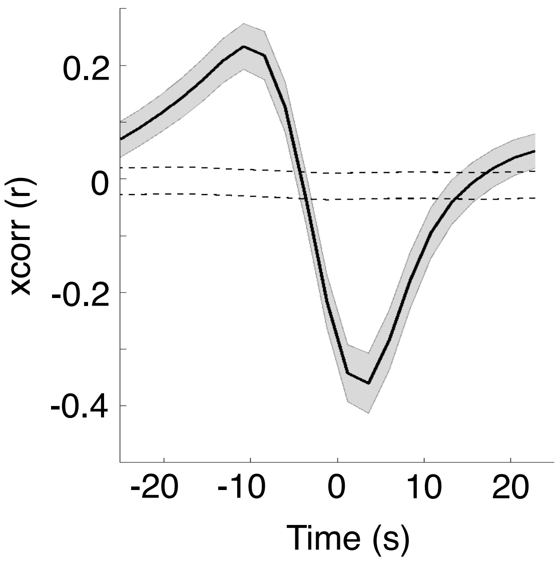


CSF time course was extracted from the BOLD-fMRI data by averaging the signals of voxels within the fourth ventricle (segmented using the FreeSurfer "aseg" atlas). Consistent with previous reports[9], we observed a strong anti-correlation between CSF and the global fMRI signal, supporting a dominant inflow contrast despite the low imaging TR employed in the study. Positive lags indicate delayed CSF than BOLD dynamics. Mean and standard errors across subjects are shown (*N*=23). Gray dashed lines indicate 95% intervals of the null group-averaged correlations, derived nonparametrically from phase-reshuffled data.
